# Extraction of directional electron-density features from diffraction data using spherical-harmonic decomposition

**DOI:** 10.64898/2026.08.04.742922

**Authors:** Santosh Panjikar, Manfred S. Weiss, Dylan Jayatilaka

## Abstract

Directional anisotropy in electron density provides key information about chemical bonding that is not readily accessible from conventional electron-density maps.

Here, a model-independent framework is presented for decomposing experimental structure factors into angular components using spherical harmonics. Reciprocal-space projection onto spherical harmonics followed by standard Fourier synthesis yields angularly filtered density maps. The *ℓ* = 0 component captures the isotropic part of the density, while the *ℓ* = 1 components resemble *p_x_*, *p_y_* and *p_z_*–like dipolar functions that highlight directional electronic structure.

Applications to high-resolution datasets, including urea, the Gly–Ala dipeptide and a 0.97 Å*β*-lactamase structure, reveal chemically interpretable dipolar features associated with carbonyl and amide bonds, N–H interactions and aromatic *π* systems. Quantitative analysis using bond-centred sampling demonstrates stable dipolar signatures that remain detectable under moderate resolution truncation.

These results establish spherical-harmonic angular decomposition as a practical framework for extracting directional electronic information from crystallographic electron-density maps.

**Synopsis:** Angular decomposition of experimental structure factors reveals dipolar anisotropy and directional electron-density features that are directly meaningful for chemical interpretation.

## 1 Introduction

Electron density derived from X-ray diffraction provides the primary link between crystallographic structure and chemical interpretation. Directional variations within the electron density encode information about bond polarity, lone-pair orientation, and intermolecular interactions, yet these features can be difficult to distinguish in conventional electron-density maps, which are dominated by isotropic contributions.

High-resolution charge-density X-ray diffraction studies of small molecules have demonstrated that experimental electron densities deviate from spherical atomic models and exhibit chemically meaningful aspherical features associated with bonding and lone-pair structure (Koritsanszky & Coppens, 2001).

These features are commonly analysed using multipolar models, particularly within the Hansen–Coppens formalism (Hansen & Coppens, 1978; Coppens, 1997), and have provided detailed insight into chemical reactivity, intermolecular interactions, and material properties (Stalke, 2011; Grabowsky *et al*., 2017; Tolborg & Iversen, 2019).

In macromolecular crystallography, however, such directional features are more difficult to resolve because typical diffraction data are obtained at lower spatial resolution, where fine angular structure becomes progressively attenuated.

Advances in synchrotron instrumentation, automated data collection and refinement strategies have led to a steady increase in the number of macromolecular structures determined at sub-Å and near-atomic resolution (Petrova & Podjarny, 2004). The growing number of macromolecular structures refined at better than 1.2 Å resolution illustrates these technical developments (Fig. 1).

**Figure 1:**
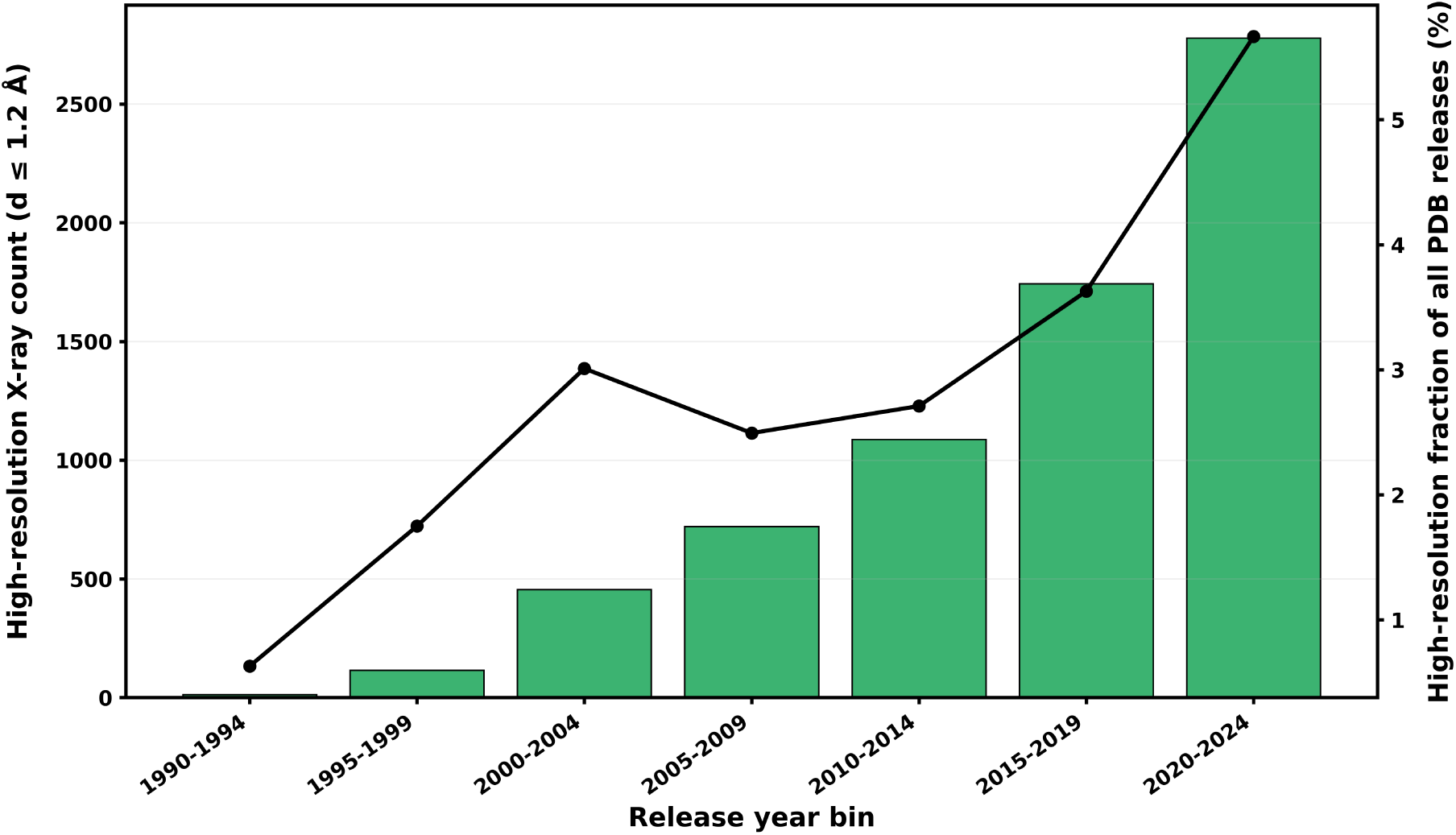
Growth of high-resolution X-ray crystal structures in the Protein Data Bank. Bars show the number of PDB entries released in each 5-year bin that were determined by X-ray diffraction at *d* ≤ 1.2 Å (left axis). The black line (right axis) shows the same quantity expressed as a percentage of all PDB releases in the corresponding 5-year bin. Together, the plot highlights both the increase in absolute numbers of sub-1.2 Å structures and their rising share of total PDB deposits in recent years. *et al*., 2005; Guillot *et al*., 2008)), as demonstrated in ultra-high-resolution studies of systems including aldose reductase (Muzet *et al*., 2003), crambin (Jelsch *et al*., 2000), trypsin (Schmidt *et al*., 2003), scorpion toxin (Pichon-Pesme *et al*., 2004), and cholesterol oxidase (Zarychta *et al*., 2015). However, such refinements typically require a substantial increase in the number of adjustable parameters and careful use of constraints and restraints, particularly for large macromolecular systems. These practical considerations have limited the routine application of multipolar approaches in macromolecular crystallography.

Despite these advances, conventional macromolecular refinement strategies continue to rely on simplified atomic models, and directional features associated with chemical bonding remain challenging to interpret directly from conventional electron-density maps. These limitations motivate approaches that extract chemically meaningful directional information from experimental electron density.

In routine macromolecular refinement, electron density is commonly interpreted within the Independent Atom Model (IAM), in which atoms are represented by spherically averaged densities. Although highly successful for structure determination, this approximation does not explicitly represent directional features of electron density that may contain chemically or functionally significant information (Urzhumtsev & Lunin, 2019).

Multipolar models based on the Hansen–Coppens formalism (Hansen & Coppens, 1978; Coppens, 1997) have enabled recovery of bonding features in macromolecules (see, for example, (Jelsch

These considerations motivate alternative strategies that retain the simplicity of conventional crystallographic refinement while enabling extraction of directional information directly from experimental data. Spherical harmonic functions provide a natural mathematical basis for separating isotropic and anisotropic components of three-dimensional density distributions. When applied in reciprocal space, spherical-harmonic projections enable angular decomposition of experimentally measured structure factors without modifying the underlying crystallographic model.

The lowest-order anisotropic term (*ℓ* = 1) of the spherical-harmonic expansion corresponds to dipolar structure and represents the dominant directional component expected in many chemically relevant environments. Dipolar anisotropy arises naturally from asymmetric charge distributions associated with polarised covalent bonds, hydrogen-bonding interactions and directional electrostatic environments. Focusing on the *ℓ* = 1 contribution therefore provides a physically interpretable representation of directional density while maintaining numerical robustness.

In the present work, spherical-harmonic angular decomposition is applied directly to experimentally derived structure factors to obtain dipolar (*ℓ* = 1) electron-density components. The procedure is implemented by applying spherical-harmonic projectors in reciprocal space followed by standard Fourier synthesis, yielding three orthogonal dipolar component maps aligned with the crystallographic *x*, *y* and *z* directions. A schematic overview of the method is shown in Fig. 2. All dipolar maps presented in this work are expressed in the global crystallographic reference frame.

**Figure 2:**
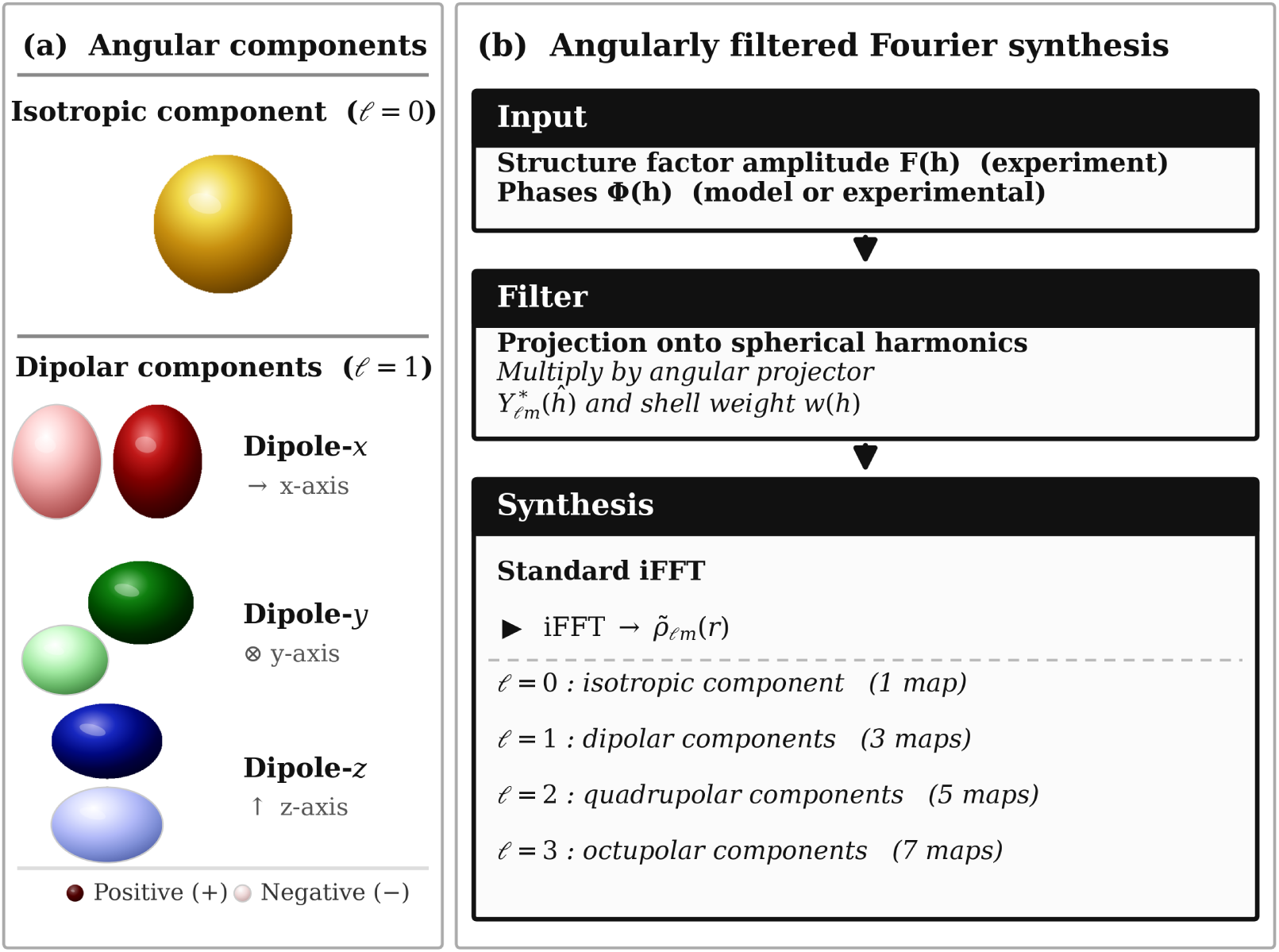
Angular decomposition of crystallographic electron density and angularly filtered Fourier synthesis. (a) Real-space angular components used in this work. The isotropic component (*ℓ* = 0) is shown as a reduced sphere, representing angular averaging rather than atomic spherical symmetry. The three dipolar components (*ℓ* = 1) correspond to charge polarisation along the *x*, *y* and *z* directions. Dark and light lobes indicate positive and negative isosurfaces. (b) Schematic of the angularly filtered Fourier synthesis. Experimental structure-factor amplitudes and phases are multiplied by spherical-harmonic projectors and shell weights prior to inverse Fourier transformation (iFFT), yielding angular-momentum-resolved real-space maps *ρ_ℓm_*(**r**), where *ρ_ℓm_*(**r**) denotes the real-space electron-density component corresponding to angular order (*ℓ, m*). Higher-order components (*ℓ* ≥ 2) are obtained in an analogous manner.

**Figure 3:**
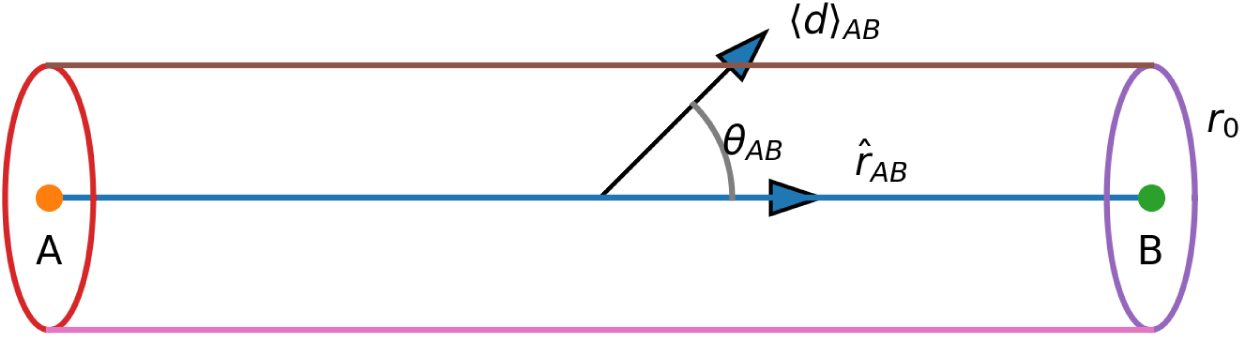
Bond-centred sampling geometry and dipolar anisotropy. For a bond *A*–*B*, density fields are sampled within a cylinder of radius *r*_0_ aligned with the bond-direction unit vector ^**r***_AB_*. Volume-averaging of the *ℓ* = 1 dipolar field within the cylinder yields the mean dipole vector ⟨**d**⟩*_AB_*. The alignment angle *θ_AB_* between ⟨**d**⟩*_AB_* and ^**r***_AB_* quantifies directional dipolar alignment.

The resulting dipolar maps are evaluated using both high-resolution small-molecule datasets and macromolecular crystallographic data. Bond-centred dipole descriptors are introduced to quantify directional density relative to chemically defined reference frames, enabling systematic analysis of bond polarity and hydrogen-bond alignment.

Numerical validation demonstrates preservation of symmetry, rotational behaviour and orthogonality between harmonic orders, confirming the stability of the recovered angular observables.

Together, these developments establish spherical-harmonic angular decomposition as a practical framework for extracting directional information from conventional crystallographic data.

## 2 Notation

Throughout this work, real-space positions are expressed in fractional unit-cell coordinates, and reciprocal-space quantities are expressed using Miller indices.

Real-space positions are denoted by ***r***, with the Fourier phase factor written as

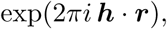

where ***h*** = (*h, k, l*) denotes Miller indices.

Reciprocal-space vectors are denoted by ***k*** (Å^−1^), with magnitude *k* = ∥*k*∥. The reciprocal-space magnitude is related to the real-space resolution *d* by

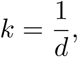

where *d* denotes the interplanar spacing associated with the reflection, so that *k* represents the magnitude of the reciprocal-space vector on the resolution shell.

Angular coordinates on the unit sphere are denoted by

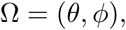

where *θ* is the polar angle measured from the positive *z* axis, and *ϕ* is the azimuthal angle measured in the *x*–*y* plane about the *z* axis.

Spherical harmonics are denoted by *Y_ℓm_*(Ω) and defined using the Condon–Shortley phase convention, where

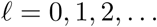

denotes the angular degree (multipole order) and

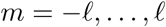

denotes the azimuthal index.

In particular, *ℓ* = 0 corresponds to isotropic terms, *ℓ* = 1 corresponds to dipolar terms, and higher values of *ℓ* represent progressively higher-order angular structure.

Spherical Bessel functions of the first kind are denoted by *j_ℓ_*.

Electron-density fields are denoted by

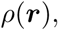

representing the conventional electron density obtained by Fourier synthesis of structure factors.

Angularly filtered density components are denoted by

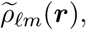

where the tilde indicates reconstruction from structure factors that have been modified by spherical-harmonic weighting in reciprocal space.

For *ℓ* = 0,

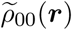

corresponds to the isotropic component, whereas higher-order terms (*ℓ* ≥ 1) represent anisotropic contributions.

*P_ℓ_*(*k*) Reciprocal-space angular power spectrum evaluated on shells of radius *k*.

*P_ℓ_*(***r***) Real-space angular power, representing the local magnitude of multipolar anisotropy.

The reciprocal-space angular power spectrum *P_ℓ_*(*k*) characterises the distribution of angular structure as a function of resolution in reciprocal space, whereas the real-space power *P_ℓ_*(***r***) provides a spatially resolved measure of the magnitude of anisotropy in real space. The dipolar case *P*_1_(***r***) is defined explicitly in Eq. (10).

These conventions are used consistently throughout the Mathematical Framework and Appendix A, unless stated otherwise.

## 3 Mathematical framework

The angular decomposition employed in this work is formulated as a reciprocal-space projection procedure applied directly to crystallographic structure factors. Each reflection is projected onto spherical-harmonic angular channels, followed by conventional inverse Fourier synthesis to obtain angularly filtered real-space density components.

The theoretical foundation of this approach is provided by the spherical Fourier–Bessel representation of electron density, derived from the Rayleigh expansion of plane waves. In the continuous limit, this formulation expresses the angular structure of reciprocal-space density through spherical-harmonic projections at fixed reciprocal radius. For completeness, this continuous theoretical formulation and its mathematical properties are presented in Appendix A.

In practical crystallographic applications, structure factors are available only at discrete reciprocal-lattice points. The implementation adopted here therefore introduces a discrete shell-wise approximation to the continuous spherical formulation. Radial integration is replaced by finite reciprocal-space shells, allowing angular projections to be evaluated using standard FFT-based Fourier synthesis, without explicit evaluation of spherical Bessel functions.

Thus, Appendix A establishes the continuous mathematical framework, while the present section develops its discrete realization suitable for experimental crystallographic data. The derivation below therefore begins from the conventional Fourier representation of electron density and introduces the angular projection step in a form directly compatible with crystallographic structure factors. Numerical validation of this discrete implementation is described in Supplementary Section S1.

The present approach does not constitute an orbital decomposition and does not treat the electron density as a wavefunction. Instead, it decomposes the angular dependence of the crystallographic electron-density field itself by projecting its reciprocal-space representation onto spherical-harmonic channels. The resulting components therefore quantify directional anisotropy of the density rather than amplitudes of underlying electronic orbitals.

The angular decomposition employed in this work is formulated as a reciprocal-space projection applied to crystallographic structure factors. The derivation begins from the standard Fourier representation of the electron density within the crystallographic unit cell.

### Starting point: conventional Fourier synthesis

Electron density in fractional coordinates ***r*** is reconstructed from structure factors using the standard inverse Fourier relation,

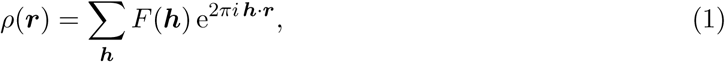

where ***h*** = (*h, k, l*) denotes Miller indices, ***r*** fractional coordinates within the unit cell, and

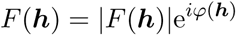

are the complex structure factors with measured amplitudes |*F* (***h***)| and phases *φ*(***h***).

Equation (1) expresses the electron density as a superposition of plane waves associated with reciprocal-lattice vectors.

#### Reciprocal-space representation in spherical coordinates

To analyse directional variation in reciprocal space, reciprocal-lattice vectors are expressed in spherical coordinates. Each reciprocal vector ***k_h_***derived from Miller indices ***h*** is characterized by its magnitude

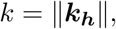

and its direction specified by the unit vector

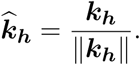

The angular coordinates Ω = (*θ, ϕ*) are defined with respect to a fixed Cartesian reference frame. For a reciprocal vector ***k_h_***= (*k_x_, k_y_, k_z_*), the polar and azimuthal angles are given by

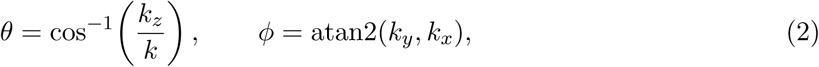

where *θ* ∈ [0*, π*] denotes the polar angle measured from the positive *z* axis, and *ϕ* ∈ (−*π, π*] denotes the azimuthal angle in the *xy* plane.

This transformation separates reciprocal-space variables into radial and angular components. The radial coordinate *k* determines spatial frequency, while angular coordinates (*θ, ϕ*) describe directional dependence.

Such separation provides the natural coordinate system for projection onto spherical harmonics *Y_ℓm_*(***k***), which form an orthonormal basis on the unit sphere.

#### Angular projection of reciprocal-space structure factors

To extract directional information, the reciprocal-space representation is projected onto spherical-harmonic angular basis functions. Using the spherical representation defined above, angular variation is described using spherical harmonics *Y_ℓm_*(***k***), which form an orthonormal basis on the unit sphere.

Directional components of the electron density are obtained by multiplying each structure factor by the angular projector 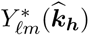. Applying this projection to Eq. (1) yields the angularly filtered density

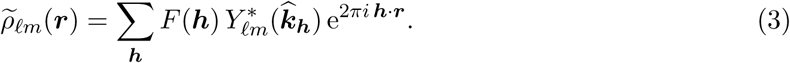

Equation (3) represents a direct angular projection of reciprocal-space structure factors onto spherical-harmonic components.

#### Discrete shell normalization

In the continuous spherical Fourier–Bessel formulation (Appendix A), angular components are obtained through integration over reciprocal-space shells with volume element *k*^2^ d*k* dΩ.

Because crystallographic data are available only at discrete reciprocal-lattice points, this radial integration is approximated by partitioning reciprocal space into finite shells of thickness Δ*k*. Reflections belonging to shell *s* are assigned a normalization factor that approximates the corresponding shell volume.

Including this normalization leads to the discrete form used throughout this work,

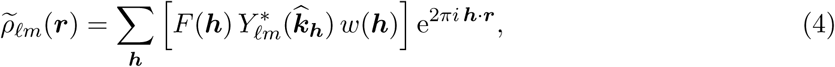

which corresponds to the angularly filtered Fourier synthesis employed in the present method. The weighting factor *w*(***h***) provides a discrete approximation to the spherical integration measure *k*^2^ d*k*, ensuring consistent normalization across reciprocal-space shells.

This discrete expression follows from the spherical Fourier–Bessel expansion of plane waves derived in Appendix A, where the continuous formulation is presented in full.

Grouping reflections into shells is therefore not required for the Fourier synthesis itself, but provides a practical approximation to the continuous spherical formulation in which angular projections are defined on surfaces of constant reciprocal-space radius. Without shell-based normalization, different regions of reciprocal space would contribute unevenly, leading to biased angular power estimates.

#### Role of the complex conjugate

The complex conjugate 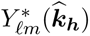 appears in Eq. (4) as a consequence of the standard inner-product definition used in spherical-harmonic analysis. In the continuous formulation, angular coefficients are obtained by projecting a function *f* (***k***^^^) onto spherical harmonics using

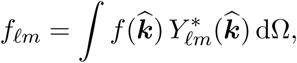

which ensures orthogonality under the normalization condition

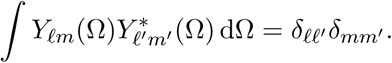

The use of the complex conjugate therefore guarantees that each angular component (*ℓ, m*) is extracted consistently according to the orthogonality properties of the spherical-harmonic basis.

#### Shell-weight normalization

In the continuous spherical Fourier–Bessel formulation, angular components of the reciprocal-space representation are obtained through integration over spherical shells with volume element *k*^2^ d*k* dΩ. Integration over the angular variables dΩ yields the radial measure 4*πk*^2^ d*k*, which defines the contribution of each reciprocal-space shell to the total integral.

Because crystallographic data are available only at discrete reciprocal-lattice points, this continuous radial integration is approximated by partitioning reciprocal space into finite shells bounded by [*κ_s_, κ_s_*_+1_). The corresponding shell volume is therefore

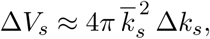

where 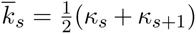 is the mean reciprocal-space radius of shell *s*, and Δ*k_s_* = *κ_s_*_+1_ − *κ_s_* is the shell thickness.

To obtain a discrete approximation, this shell volume is distributed uniformly among the reflections that belong to the shell. If *N_s_* denotes the number of observed reflections in shell *s*, each reflection is assigned a weight equal to the shell volume divided by the number of contributing reflections.

The resulting shell-weight factor is defined as

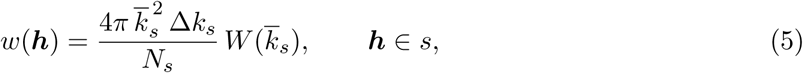

where *W* (*k*) is an optional radial tapering function. In its simplest form, *W* (*k*) = 1 for all reflections. Optional smooth tapering functions (e.g. cosine or Kaiser windows) may be applied near the maximum reciprocal-space radius to reduce truncation artefacts associated with finite resolution limits.

This normalization ensures that each reciprocal-space shell contributes proportionally to its spherical volume, providing a discrete Riemann approximation to the continuous radial integral. Importantly, division by *N_s_* distributes the shell volume only among observed reflections, allowing the normalization to remain well defined even in the presence of incomplete reciprocal-space sampling.

#### Optional radial tapering

In its simplest implementation, no additional radial modification is applied, and the tapering function is defined as

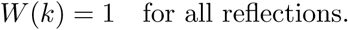

Under this condition, Eq. (5) reduces to a pure shell-volume normalization, and the angular decomposition is performed directly on the measured structure factors.

Optional smooth tapering functions *W* (*k*) may be introduced near the maximum reciprocal-space radius *k*_max_ to reduce truncation artefacts associated with finite-resolution limits. In the continuous spherical formulation, abrupt termination of reciprocal-space data produces oscillatory artefacts (Gibbs-type ringing) in the reconstructed density. Applying a smooth radial taper gradually suppresses contributions near *k*_max_, thereby improving numerical stability and reducing high-frequency noise.

Typical choices include cosine or Kaiser window functions (Harris, 1978; Kaiser, 1974), although the precise functional form is not critical. Use of tapering is therefore optional and serves only as a numerical regularization strategy; it is not required for the validity of the angular decomposition itself.

In addition to reducing truncation artefacts, radial tapering may also improve numerical stability when the highest-resolution shells are incompletely sampled. Gradual attenuation of contributions near *k*_max_ reduces the influence of sparsely populated shells, thereby stabilizing angular power estimates.

#### Incomplete high-resolution sampling

In practical crystallographic datasets, the highest-resolution reciprocal-space shells are often incompletely sampled due to experimental limitations or anisotropic data collection. Because angular decomposition involves normalization of contributions within reciprocal-space shells, sparsely populated outer shells may introduce increased statistical variability in the recovered angular components, particularly for higher-order terms.

The shell-weight normalization defined in Eq. (5) remains well defined under incomplete sampling, since weights are distributed only among observed reflections. However, when the outermost shells contain very few reflections, their contribution to the reconstructed angular components may become noisy. Optional radial tapering functions *W* (*k*) provide a practical means to gradually attenuate contributions near the maximum reciprocal-space radius *k*_max_, thereby reducing sensitivity to sparsely sampled high-resolution regions.

The numerical behaviour of the present implementation under incomplete reciprocal-space sampling was evaluated using synthetic incompleteness tests, including random and directional removal of reflections (Supplementary Section S1). These tests confirm that the shell-weight normalization remains stable and that optional tapering improves robustness when outer-shell completeness is reduced.

### 3.1 Real dipolar components and truncation

The angular components introduced above are expressed in the complex spherical-harmonic basis. For visualization and chemical interpretation, these complex components were transformed into real (Cartesian) dipolar maps using standard linear combinations of the *ℓ* = 1 harmonic components.

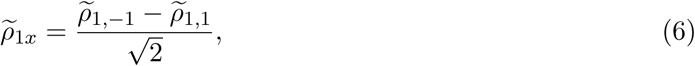

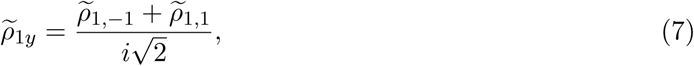

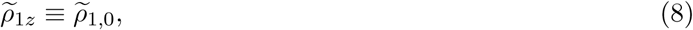

yielding three orthogonal dipolar components aligned with the crystallographic *x*, *y*, and *z* directions (Fig. 2). These real components together form a vector dipolar field

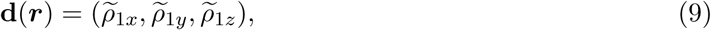

which represents the local dipolar anisotropy of the electron density.

This vector field transforms under rotation in the same manner as a classical dipole, providing a direct geometrical interpretation of directional density asymmetry.

Each dipolar component retains the same numerical map-value units as the underlying density e Å^−3^, but represents a signed anisotropic contribution rather than a total electron density. Consequently, the *ℓ* = 1 maps exhibit characteristic positive and negative lobes that describe the directional distribution of density around each spatial location.

In addition to the vector field representation, a rotationally invariant scalar measure of dipolar strength can be defined as the dipolar power

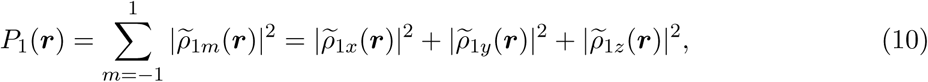

which provides a coordinate-independent measure of the magnitude of dipolar anisotropy at each spatial location.

Because this quantity depends only on squared magnitudes of the components, it remains invariant under coordinate rotation. Under rotation of the coordinate system, the individual coefficients *ρ*_1_*_m_* are redistributed among different values of *m*, but their total power *P*_1_ remains unchanged.

#### Rotational invariance of dipolar power

The rotational invariance of *P*_1_(***r***) follows directly from the transformation properties of spherical harmonics.

Under rotation of the coordinate system, the *ℓ* = 1 coefficients transform through a unitary rotation

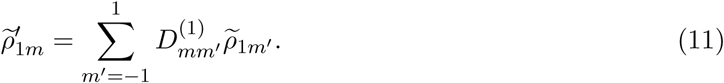

The dipolar power in the rotated frame is

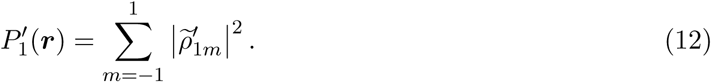

Substituting Eq. (11) yields

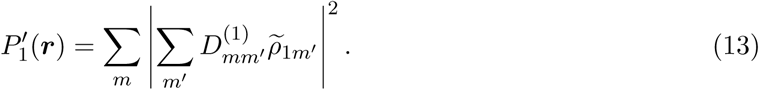

Because the rotation matrix 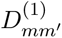 is unitary,

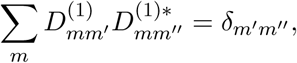

it follows directly that

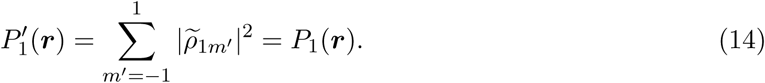

Thus, the dipolar power is invariant under coordinate rotation, even though individual coefficients *ρ*_1*m*_ are redistributed among different values of *m*.

This mathematical invariance forms the theoretical basis for the numerical rotational tests described in Supplementary Section S1.

The isotropic component (*ℓ* = 0) is recovered naturally from the same shell-wise filtering framework. Apart from the constant normalization factor *Y*_00_ = 1*/*^√^4*π*, this component corresponds directly to the conventional crystallographic electron density.

In the present work, primary analyses focus on the lowest-order anisotropic contribution (*ℓ* = 1), which provides the simplest physically interpretable representation of directional density variations.

Higher-order components (*ℓ* ≥ 2) were evaluated in validation studies (Supplementary Section S1) to assess numerical convergence and orthogonality of the harmonic expansion. Although these higher-order terms encode additional angular structure, the present manuscript focuses on the dipolar (*ℓ* = 1) contribution, while leaving detailed analysis of higher-order components to future investigations.

These definitions establish the practical real-space representation of dipolar anisotropy. This representation is used throughout the analysis.

### 3.2 Physical interpretation of harmonic orders

Each harmonic order (*ℓ*) corresponds to a characteristic angular symmetry pattern, providing a direct link between mathematical decomposition and chemically meaningful directional structure.

The isotropic component (*ℓ* = 0) represents the spherically symmetric component of the electron density and corresponds to the isotropic part of the conventional electron-density distribution obtained from standard Fourier synthesis.

The first anisotropic order (*ℓ* = 1) corresponds to dipolar structure and represents the simplest directional asymmetry in the density. Dipolar features arise naturally from asymmetric charge distributions commonly associated with polarized covalent bonds, hydrogen-bonding interactions, directional electrostatic environments.

In these situations, electron density is displaced preferentially along a specific direction, producing the characteristic paired lobes patterns observed in dipolar-filtered maps.

Higher-order harmonics (*ℓ* ≥ 2) represent progressively more complex angular patterns, including quadrupolar (two-lobed or two-centre) symmetry, multi-centre bonding environments, and higher-order multipolar features such as those associated with aromatic *π*-electron systems or extended charge distributions.

Although these higher-order terms encode additional angular detail, the dipolar (*ℓ* = 1) contribution provides the most direct and chemically interpretable description of directional polarisation effects within the scope of the present work.

Together, these harmonic orders form a hierarchical description of angular structure, with increasing *ℓ* capturing progressively finer directional detail in the electron density.

Although the present analyses are expressed in the global crystallographic reference frame, the same dipolar representation can be transformed into chemically defined local coordinate systems, such as bond-centred or atom-centred frames. Such local transformations enable quantitative comparison of directional density relative to specific molecular geometries, including bond directions and hydrogen-bond alignments.

Exploration of local-frame descriptors provides a natural extension of the present framework and will be addressed in future studies.

## 4 Method

### 4.1 Dataset

Small-molecule benchmark datasets were selected from high-resolution entries in the Cambridge Structural Database (CSD), chosen to provide chemically well-characterised reference systems with accurately determined hydrogen positions, minimal thermal motion, and high-resolution diffraction data. Such systems provide reliable benchmarks for evaluating directional features in electron density.

The glycine–alanine dipeptide was analysed using the dataset with CSD refcode GLYALB05 (CCDC deposition number 995876), refined against X-ray data collected at 12 K to 0.66 Å resolution using Hirshfeld atom refinement (HAR) as implemented in Tonto (Jayatilaka & Grimwood, 2001; Jayatilaka & Dittrich, 2008; Capelli *et al*., 2014). This dataset provides a well-characterised peptide-bond system with accurately determined hydrogen positions.

Urea was taken from the CSD entry UREA01 and refined against sub-Å X-ray data (0.35 Å resolution) reported by Birkedal *et al*. (Birkedal *et al*., 2004). The corresponding reflection data were obtained from the IUCr supplementary material for that study and processed using the same analysis workflow.

For the CSD datasets, observed structure-factor amplitudes together with phases calculated from the corresponding refined models were used for Fourier synthesis, consistent with standard crystallographic density reconstruction procedures. Key crystallographic parameters for all datasets are summarised in Table 1.

**Table 1:**
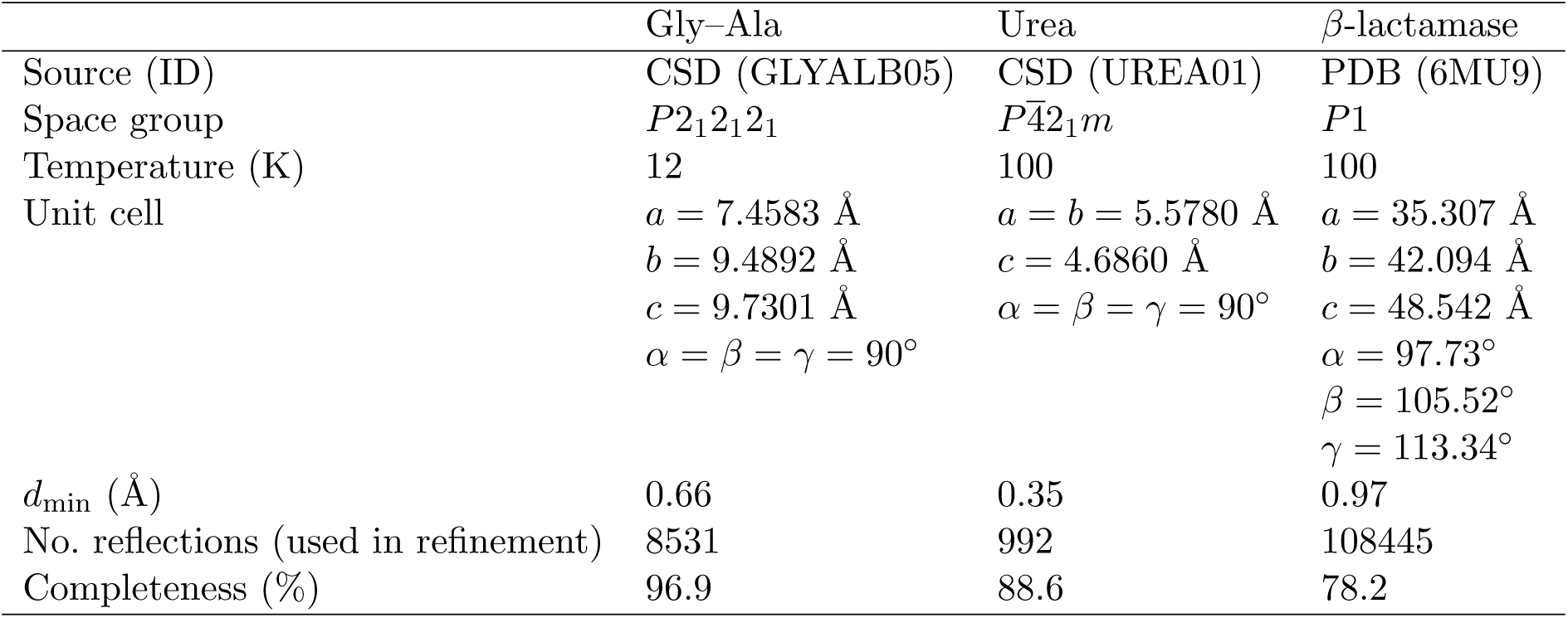
Crystallographic data used in this work. For the macromolecular MTZ dataset, completeness was computed as the fraction of unique reciprocal-ASU reflections present within the stated resolution range. For the CSD datasets, completeness values were taken from CIF metadata where available and correspond to reported unique-reflection completeness.

| | Gly-Ala | Urea | β-lactamase |
| --- | --- | --- | --- |
| Source (ID) | CSD (GLYALB05) | CSD (UREA01) | PDB (6MU9) |
| Space group | $P2_12_12_1$ | $P\bar{4}2_1m$ | $P1$ |
| Temperature (K) | 12 | 100 | 100 |
| Unit cell | $a = 7.4583 \text{ \AA}$<br>$b = 9.4892 \text{ \AA}$<br>$c = 9.7301 \text{ \AA}$<br>$\alpha = \beta = \gamma = 90^\circ$ | $a = b = 5.5780 \text{ \AA}$<br>$c = 4.6860 \text{ \AA}$<br>$\alpha = \beta = \gamma = 90^\circ$ | $a = 35.307 \text{ \AA}$<br>$b = 42.094 \text{ \AA}$<br>$c = 48.542 \text{ \AA}$<br>$\alpha = 97.73^\circ$<br>$\beta = 105.52^\circ$<br>$\gamma = 113.34^\circ$ |
| $d_{\min}$ (Å) | 0.66 | 0.35 | 0.97 |
| No. reflections (used in refinement) | 8531 | 992 | 108445 |
| Completeness (%) | 96.9 | 88.6 | 78.2 |

Macromolecular analyses were performed using the high-resolution *β*-lactamase structure (PDB entry 6MU9), refined at 0.97 Å resolution by the original authors using REFMAC5 (Murshudov *et al*., 1997) within the Independent Atom Model (IAM).

Complete structure-factor amplitudes and phases (*F* and *ϕ*) were available for this dataset and were used directly for Fourier synthesis. This structure has previously been used to investigate peptide-bond chemistry and protonation-related difference-density features, providing a well-characterised macromolecular reference system for the present analysis (Panjikar & Weiss, 2025).

Together, these datasets span sub-Å to near-Å resolution, providing chemically well-resolved reference systems for validating the extraction and interpretation of directional electron-density features across both small-molecule and macromolecular regimes.

Although the underlying datasets differ in crystal system, resolution, and reflection sampling, conversion of the small-molecule data to MTZ format enabled all datasets to be processed using a common Fourier-synthesis and spherical-harmonic projection pipeline, ensuring methodological consistency across all analyses.

### 4.2 Reflection preparation: phases, resolution selection, symmetry and Friedel expansion

Structure-factor amplitudes |*F* (***h***)| and corresponding phases *φ*(***h***) were obtained from refined crystallographic models using standard software (e.g. CCP4 suite programs). No modification of the atomic model or refinement target was performed during the angular decomposition.

Only reflections within the stated resolution range were included. The reciprocal-space resolution was defined as *d* = 1/ ∥*k*∥***k_h_***, where ***k_h_*** is the reciprocal vector associated with Miller indices ***h***.

Reflections with missing amplitudes or undefined phases were excluded.

To provide uniform reciprocal-lattice sampling required for angular filtering, input reflection sets were expanded to space group *P* 1 prior to map calculation. For datasets deposited in space groups other than *P* 1, symmetry-related reflections were generated by applying the crystallographic symmetry operations of the deposited space group.

Where anomalous data were present, Friedel mates were included explicitly by Friedel expansion, ensuring that both ***h*** and −***h*** reflections were available. This step provides symmetric directional sampling in reciprocal space, which is required for consistent angular projection onto spherical harmonics. No modification of measured amplitudes or phases was performed; when anomalous differences were present, both members of each Friedel pair were retained. Angular filtering was applied after symmetry and Friedel expansions, and all angularly filtered maps were computed by inverse Fourier transformation on a Cartesian grid in *P* 1. No local site-symmetry constraints were imposed, and all angular components were evaluated in the global crystallographic reference frame.

### 4.3 Angular basis, dipolar channels, and truncation

For each reflection ***h***, angular projectors *Y* ^∗^ (***k_h_***) were evaluated using the complex spherical-harmonic basis. The unit direction ***k_h_*** was computed directly from the reciprocal-lattice geometry using the unit-cell metric and Miller indices.

The angular expansion was truncated at *ℓ* = 1, corresponding to the lowest-order aspherical contribution to the electron density. The *ℓ* = 1 components represent dipolar anisotropy and therefore provide the lowest-order description of directional charge redistribution relative to the isotropic density. Three complex dipolar channels (*ℓ, m*) = (1, −1), (1, 0), (1, 1) were evaluated independently. Real dipolar components aligned with the Cartesian axes were constructed by the standard linear combinations defined in Eqs. (6)–(8), yielding orthogonal *x*-, *y*- and *z*-dipolar maps corresponding to the Cartesian components of the *ℓ* = 1 field (Fig. 2).

At typical macromolecular resolutions, dipolar (*ℓ* = 1) anisotropy represents the lowest-order deviation from spherical symmetry that remains robust under finite sampling, making it the most reliable directional descriptor accessible from conventional diffraction data.

All calculations were performed using data truncated at the stated high-resolution limit. No extrapolation beyond the measured resolution was attempted. Higher-order terms (*ℓ* ≥ 2) can be obtained using the same angular filtering and synthesis procedure, but were not included here in order to focus on the dominant dipolar contribution and to avoid overinterpretation of higher-order components at typical macromolecular resolutions.

### 4.4 Reciprocal-space discretisation, shell weights, and filtered Fourier synthesis

Angular filtering was applied in reciprocal space prior to inverse Fourier transformation. After expansion of the reflection set to *P* 1 (including Friedel mates when present), reflections were grouped into resolution shells defined by uniform bins in reciprocal-space radius *k* = ∥***k_h_***∥ = 1*/d**_h_***.

The reciprocal-space range between *k*_min_ and *k*_max_ = 1*/d*_min_ was divided into *N*_shell_ = 80 uniform radial shells, giving a constant shell thickness

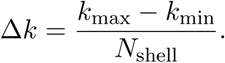

The number of shells (80) was selected to provide adequate radial sampling while maintaining sufficient reflections per shell; tests using alternative shell counts showed negligible changes in recovered low-order angular components.

For each shell *s* with boundaries [*κ_s_, κ_s_*_+1_), the mean radius *k_s_* = <u>^1^</u> (*κ_s_* + *κ_s_*_+1_), the shell thickness Δ*k_s_* = *κ_s_*_+1_ − *κ_s_*, and the number of reciprocal-lattice samples (reflections) in the shell, *N_s_*, were evaluated. Here *N_s_* denotes the number of reflections in shell *s* after symmetry expansion; it does not represent crystallographic multiplicity.

Shell-wise normalisation weights were computed according to Eq. (5). No radial tapering was applied (i.e. uniform radial weighting), and therefore *W* (*k_s_*) = 1 for all shells. This choice implements a discrete Riemann approximation to the continuum spherical measure *k*^2^ d*k* dΩ. The factor 4*π k*^2^*_s_* Δ*k_s_* represents the reciprocal-space shell volume element, while division by *N_s_* yields a shell-wise average over the discrete sampling of the angular surface *S*^2^.

Angularly filtered maps were computed using Eq. (4) by multiplying each complex structure-factor coefficient |*F* (***h***)|e*^iφ^*^(***h***)^ by the spherical-harmonic projector *Y* ^∗^ (***k_h_***) and the corresponding shell weight *w*(***h***). The filtered coefficients were then transformed to real space using an inverse discrete Fourier transform (FFT) on a regular Cartesian grid in *P* 1.

The real-space grid spacing was set to *d*_min_*/*3, corresponding to approximately three grid points per highest-resolution spacing, a commonly used criterion for adequate Fourier sampling. This value provides adequate sampling of Fourier features while avoiding unnecessary oversampling. Grid-refinement tests showed that the recovered low-order angular observables, including the *ℓ* = 1 dipolar components, were insensitive to further grid refinement (Supplementary Section S1).

All maps were evaluated in *P* 1 and interpreted in the global crystallographic reference frame.

### 4.5 Map sampling, masking and visualisation

Unless stated otherwise, maps were computed over the full unit cell in space group *P* 1, without restricting the calculation to specific real-space regions. In particular, no spatial masking or region-of-interest selection was applied during map synthesis, although local regions were inspected visually after map calculation. All spherical-harmonic–filtered maps were therefore evaluated across the entire unit cell.

For visualisation and qualitative inspection, isotropic (*ℓ* = 0) and dipolar (*ℓ* = 1) component maps were examined both globally and locally in the vicinity of selected residues using standard crystallographic visualisation tools, primarily Coot (Emsley *et al*., 2010) and PyMOL (Schrödinger, 2015). Maps were rendered as isosurfaces or mesh contours, with positive and negative levels displayed separately to highlight directional features.

Unless stated otherwise, contour levels were set to fixed multiples of the map root-mean-square deviation, *σ*_map_, defined as the standard deviation of the map values evaluated over all grid points in the full unit cell (typically 2.5–4.0×*σ*_map_), and were kept constant across comparable maps within a given figure. For each figure, the corresponding contour values are reported in the caption, both as *n* × *σ*_map_ and, where relevant, as absolute map values reported on the map scale in e Å^−3^.

This approach avoids introducing spatial bias during map computation and maintains uniform sampling across the unit cell, thereby supporting consistent comparison of angular components between regions and datasets.

### 4.6 Quantitative dipole descriptors

#### 4.6.1 Bond-centred dipolar anisotropy descriptors

For urea (CSD refcode **UREA01**) and the glycine–alanine dipeptide (CSD refcode **GLYALB05**), structure-factor amplitudes were converted to *ℓ* = 0 (isotropic) and *ℓ* = 1 (dipolar) electron-density maps on a common Cartesian grid at the experimental resolution of the original refinements. These maps were used to define bond-centred measures of dipolar anisotropy.

For each bond *A*–*B*, the dipolar vector field **d**(**r**), defined in Eq. 9, represents the local direction and magnitude of dipolar anisotropy.

The mean dipole vector associated with bond *A*–*B* was defined by volume averaging within a cylindrical region aligned with the bond axis,

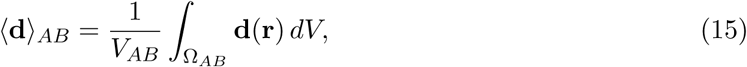

where Ω*_AB_*denotes the sampling cylinder, and *V_AB_* is its volume.

The corresponding isotropic density was evaluated over the same region,

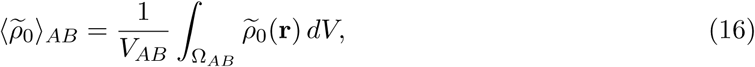

where *ρ*_0_(**r**) is the *ℓ* = 0 isotropic component.

The cylindrical sampling region Ω*_AB_* was centred on the bond axis connecting atoms *A* and *B*. The bond direction ^**r***_AB_* was defined as the unit vector pointing from the less to the more electronegative atom, using standard Pauling electronegativity values.

The cylinder extended between atoms *A* and *B* with length

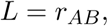

and radius

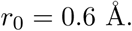

The radius r0 was selected as a compromise between spatial localisation and numerical stability, and was confirmed to yield stable descriptor values over the tested range of sampling radii (Supplementary Section S2).

Two scalar descriptors were defined. The alignment

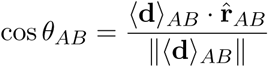

reports the orientation of the mean dipole vector relative to the bond direction. Its absolute value | cos *θ_AB_*| ranges between 0 and 1, with values approaching unity indicating strong alignment of the dipolar field with the bond direction.

A dimensionless anisotropy index

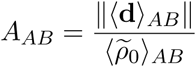

was used to describe the magnitude of the dipolar distortion relative to the local isotropic density.

These quantities were computed for all C=O, N–H and C–H bonds in urea and Gly–Ala, and grouped by bond class for statistical analysis.

### 4.7 Validation and control analyses

#### Symmetry consistency

Symmetry-related regions were compared explicitly. The corresponding dipolar features showed consistent magnitude and orientation, confirming that angular filtering after symmetry expansion preserves crystallographic symmetry relationships.

#### Friedel pairing and Hermitian consistency

Where Friedel pairs were available, angularly filtered maps were computed using both the full reflection set and explicitly Hermitian-symmetrised structure factors, *F* (**h**) and *F* ^∗^(−**h**). The resulting dipolar maps were indistinguishable within numerical precision, demonstrating that the observed *ℓ* = 1 signal arises from physically meaningful anisotropy rather than incomplete reciprocal-space sampling.

#### Resolution truncation behaviour

The stability of dipolar orientations with respect to progressive removal of high-resolution reflections was evaluated. Stable spatial localisation and orientation of dipolar features were observed for the majority of chemically defined sites. Full procedural details are provided in Supplementary Section S2.

### 4.8 Software implementation and availability

All angular decompositions and dipole descriptor calculations were performed using custom Python-based software developed for spherical-harmonic analysis of reciprocal-space data. The implementation makes use of gemmi (Wojdyr, 2021), reciprocalspaceship (Greisman *et al*., 2021), NumPy (Harris *et al*., 2020) and SciPy (Virtanen *et al*., 2020). All scripts were executed using Python 3.10 under Linux-based environments.

The software consists of command-line tools that

i. read standard MTZ or CIF reflection files,
ii. apply spherical-harmonic angular filtering and reciprocal-space shell weighting,
iii. generate angularly filtered real-space maps (*ℓ* = 0, *ℓ* = 1, and higher components),
iv. compute bond-level dipolar descriptors and associated directional metrics.

The current version of the scripts, together with example configuration files used to generate the figures in this work, is available from the corresponding author upon reasonable request. Representative input datasets and example workflows are provided to enable reproduction of the principal analyses.

## 5 Results

### 5.1 Small-molecule benchmarks: urea and glycine–alanine dipeptide

To establish a chemically unambiguous reference for the angular decomposition, the method was first applied to high-resolution small-molecule datasets. Small molecules provide a stringent benchmark because their electronic structure is well characterised and is not confounded by conformational heterogeneity or macromolecular refinement restraints.

Throughout the benchmark and protein examples, we display the *ℓ* = 1 map together with an inferred local dipole axis derived from the volume-averaged *ℓ* = 1 vector field. The *ℓ* = 1 isosurface representation localises where dipolar signal resides in real space (bond-centred versus more diffuse features), whereas the axis is computed from locally averaged *ℓ* = 1 vector components and is the quantity entering the alignment metrics (e.g. the angle *θ* relative to the C–O bond).

Because multiple dipolar contributions can overlap within the sampling region (e.g. neighbouring N–H features, lone-pair lobes, or symmetry-related density), the inferred axis should be interpreted as a *net* dipolar direction and may lie between visible lobes rather than connecting the strongest extrema.

Urea (UREA01, 0.35 Å resolution) provides a benchmark containing a strongly polarised carbonyl group within an approximately planar bonding environment, making it an ideal test case for directional anisotropy detection.Figure 4 compares the isotropic (*ℓ* = 0) and dipolar (*ℓ* = 1) components in two viewing orientations. The *Y*_00_-weighted map (Fig. 4A,C) reproduces near-spherical atomic density envelopes, whereas the combined *ℓ* = 1 dipolar component (Fig. 4B,D) isolates chemically interpretable bond-centred anisotropy.

**Figure 4:**
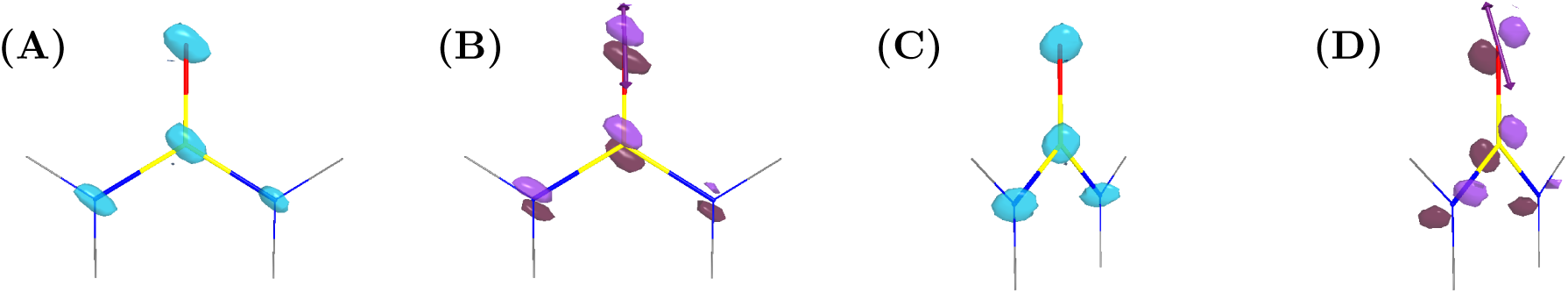
Angular decomposition of experimental electron density in urea. (A,C) Isotropic component (*ℓ* = 0; cyan isosurfaces) shown in two viewing orientations; panel C is obtained by rotating panel A by approximately 60^◦^ about the crystallographic *y* axis.(B,D) Dipolar component (*ℓ* = 1; paired positive/negative isosurfaces) shown in the same orientations; panel D corresponds to the rotated view of panel B. In (D), the dipole axis inferred from the locally averaged *ℓ* = 1 vector field is indicated by an arrow. Isosurfaces are contoured at symmetric positive and negative levels: for urea, ±23.421 e Å^−3^ for *ℓ* = 0 and ±20.126 e Å^−3^ for *ℓ* = 1 (±3.8× map r.m.s.; purple positive, maroon negative).

The *ℓ* = 0 component represents the angularly averaged part of the local density field. Although directional angular variation is removed, this component still contains contributions from neighbouring atoms and bonding regions. As a result, the *ℓ* = 0 density preserves molecular connectivity and may appear elongated along bond directions, particularly at sub-Å resolution.

The *ℓ* = 1 component reflects dipolar anisotropy modulated by the local molecular environment. Its appearance is governed by the underlying radial density distribution and bonding geometry. The inferred dipole axis represents a net directional descriptor and does not necessarily coincide with the line connecting the strongest extrema.

This separation of isotropic and dipolar contributions illustrates how directional information that is not readily visible in conventional electron-density maps becomes explicitly localised in the *ℓ* = 1 channel.

A dominant dipolar signature is aligned approximately parallel to the C=O bond, consistent with carbonyl polarisation, and weaker but directed features are visible along the N–H bonds. The dipole axis inferred from the local *ℓ* = 1 coefficients (arrow in Fig. 4D) provides a compact geometric descriptor of this anisotropy and is consistent with the expected carbonyl-associated dipolar response.

**Interpretation of the** *ℓ* = 0 **component**

The *ℓ* = 0 component corresponds to the angularly averaged part of the local density field. Although directional angular variation is removed, this component still contains contributions from neighbouring atoms and bonding regions. As a result, the *ℓ* = 0 density preserves molecular connectivity and may appear elongated along bond directions, particularly at sub-Å resolution where bonding density is strongly resolved.

The glycine–alanine dipeptide (GLYALB05, 12 K, 0.66 Å resolution) was analysed as a representative peptide system. Dipolar (*ℓ* = 1) electron-density maps for Gly–Ala (Fig. 5) exhibit pronounced anisotropy associated with the peptide linkage. These dipolar components isolate the directional part of the density distribution and therefore provide a direct visual indicator of bond polarity, which is not readily distinguishable in conventional isotropic density maps. Paired positive and negative lobes flank each carbonyl C=O group, with density displaced towards oxygen and depleted near carbon, consistent with the known polarity of peptide bonds, with electron density shifted towards oxygen and depleted near carbon. Amide N–H bonds show weaker but directionally consistent dipoles, whereas nearby C–H bonds exhibit smaller and more isotropic behaviour.

**Figure 5:**
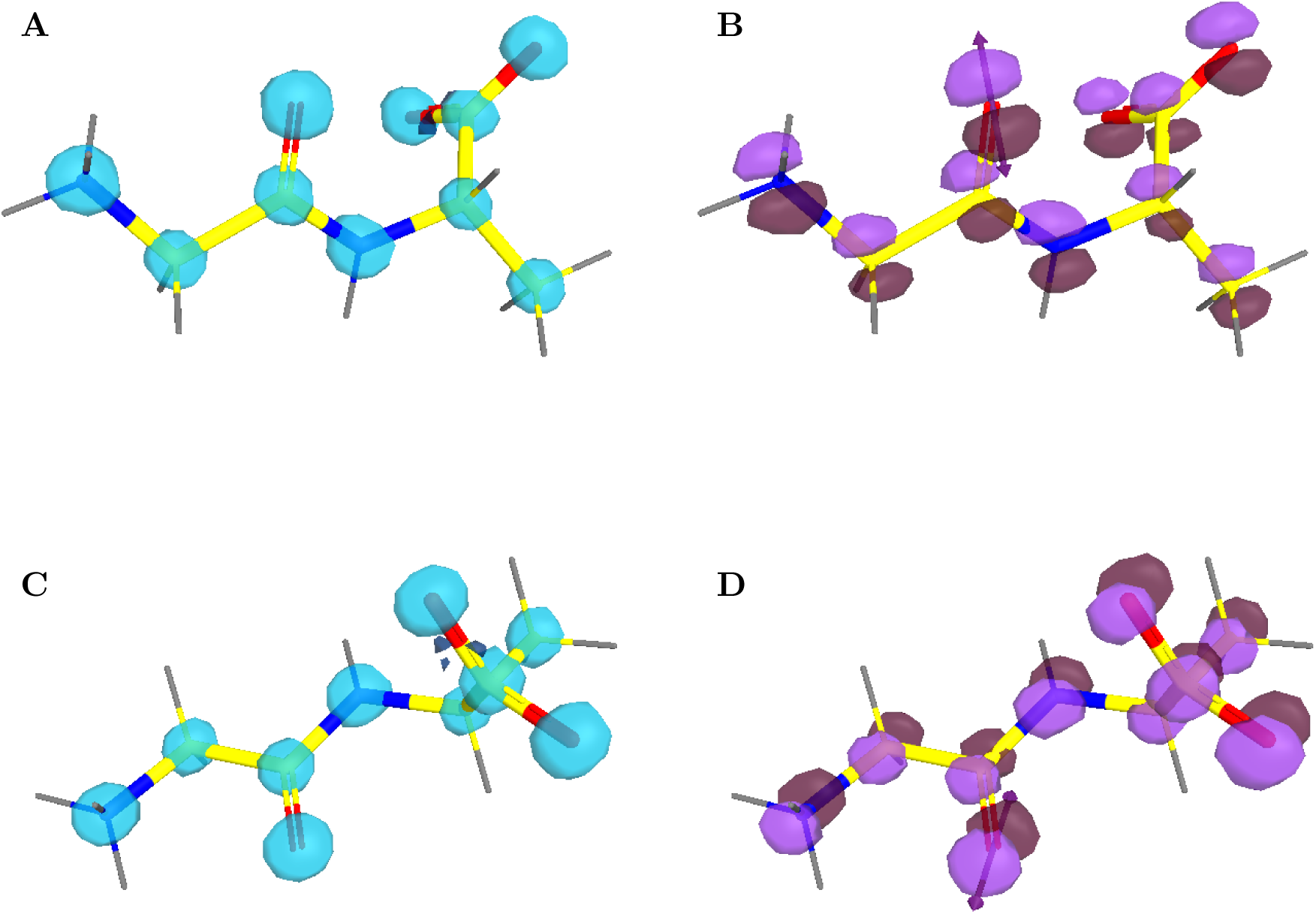
Angular decomposition of experimental electron density in the glycine–alanine dipeptide (GLYALB05) (A,C) Isotropic component (*ℓ* = 0; cyan isosurfaces) shown in two viewing orientations, reproducing the overall atomic density envelope of the dipeptide.(B,D) Dipolar component (*ℓ* = 1; paired positive and negative isosurfaces) shown in the same two orientations, highlighting bond-centred anisotropy along the peptide linkages. A dipole axis inferred from the locally averaged *ℓ* = 1 vector field at a peptide carbonyl oxygen is indicated by an arrow (most clearly visible in panel B) and provides the direction used for the quantitative alignment metrics. Panels C and D show the corresponding rotated views relative to A and B to aid interpretation of the three-dimensional dipolar features. Isosurfaces are contoured at symmetric positive and negative map levels, reported on the map scale in e Å^−3^: for Gly–Ala, ±1.738 e Å^−3^ for *ℓ* = 0 (A,C) and ±1.678 e Å^−3^ for *ℓ* = 1 (B,D), corresponding in each case to ±2.8× the map r.m.s. (purple: positive; maroon: negative).

While the isotropic (*ℓ* = 0) maps reproduce the overall atomic density envelope, the *ℓ* = 1 dipolar maps selectively highlight bond-centred directional features, revealing polarity along peptide C=O bonds and weaker anisotropy in N–H and C–H bonds.

These features are reproducible across crystallographically independent molecules, indicating that the observed dipolar signatures reflect intrinsic bonding properties rather than packing artefacts.

To quantify these observations, the bond-centred dipolar analysis described in the Methods section was applied to all C=O, N–H and C–H bonds in urea and Gly–Ala. For each bond, the absolute alignment | cos *θ_AB_*| and the anisotropy index *A_AB_* were calculated from bond-centred sampling of the *ℓ* = 1 vector field and the *ℓ* = 0 reference density. The resulting distributions are summarised in Fig. 6A for the alignment | cos *θ_AB_*| and Fig. 6B for the anisotropy index *A_AB_*.

**Figure 6:**
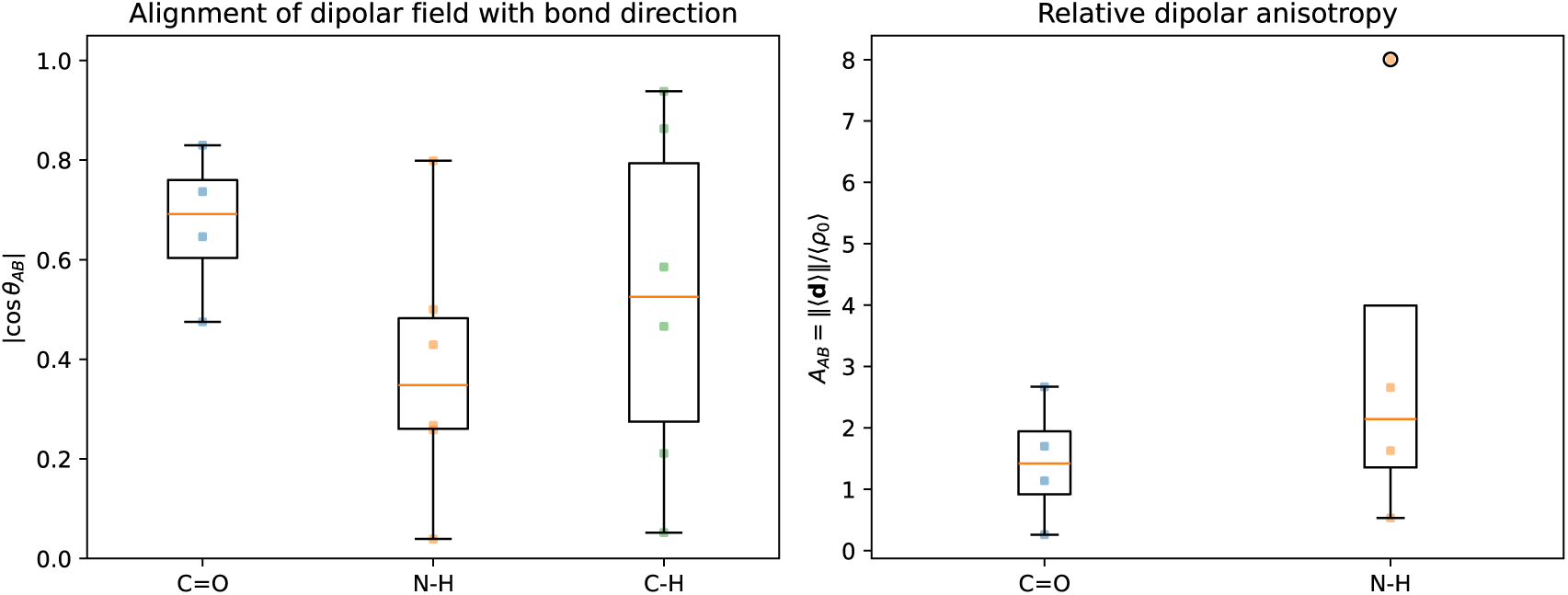
Bond-level statistics of the dipolar field for the combined urea and Gly–Ala benchmark set. (a) Distributions of the absolute alignment | cos *θ_AB_*| between the mean dipole vector and the bond direction for different bond classes, pooled across both molecules (C=O and N–H in urea; C=O, N–H and C–H in Gly–Ala). Carbonyl C=O bonds show the strongest and most coherent alignment, amide N–H bonds show intermediate alignment, and C–H bonds (present only in Gly–Ala) display the weakest and most isotropic behaviour. (b) Distributions of the anisotropy index 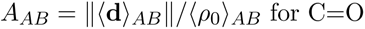 and N–H bonds. C=O bonds exhibit the largest and most stable anisotropy, with N–H bonds showing intermediate values. C–H bonds from Gly–Ala are omitted from panel (b) because the corresponding *A_AB_* values are numerically unstable, owing to the very low isotropic density at hydrogen positions.

Across the combined benchmark set, carbonyl C=O bonds exhibit the strongest and most coherent dipolar alignment, with typical values of | cos *θ_AB_*| in the range 0.6–0.8 for peptide carbonyls in Gly–Ala and approximately 0.5 for urea carbonyls, yielding an overall mean | cos *θ_AB_*| ≈ 0.7. Values approaching unity indicate strong alignment of the dipolar field with the bond axis. Amide N–H bonds display weaker and more variable alignment (mean | cos *θ_AB_*| ≈ 0.4), whereas C–H bonds show broader and more isotropic behaviour.

The dipole magnitude 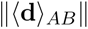 and the anisotropy index *A_AB_* follow the same qualitative hierarchy. Carbonyl C=O bonds have the largest and most stable anisotropy indices (order unity), whereas N–H bonds show intermediate values and C–H bonds often yield small or numerically unstable *A_AB_* owing to the very low isotropic density at hydrogen sites. Together, Figs. 4 and 5, and the bond-level statistics in Fig. 6, demonstrate that the *ℓ* = 1 component of the experimental density encodes chemically meaningful bond polarity and directional anisotropy in these benchmark systems, providing a calibrated reference framework for interpreting dipolar signatures in macromolecular environments.

These benchmark measurements therefore establish a calibrated reference scale for interpreting dipolar signatures in more complex macromolecular systems.

Independent numerical validation tests, including rotational invariance, translational stability, and conservation of angular power, confirm that these observations are robust with respect to coordinate choice and numerical implementation (Supplementary Section S1).

### 5.2 Dipolar anisotropy of peptide bonds in proteins

#### Extension to macromolecular peptide bonds

Having established the behaviour of the angular decomposition in chemically simple systems (small-molecule benchmarks and synthetic validation; Supplementary Section S1), the analysis was extended to peptide bonds in macromolecular structures.

Peptide bonds constitute a ubiquitous structural motif in proteins and exhibit intrinsic directional charge polarisation associated with carbonyl polarity and amide donation. These effects are not explicitly represented in conventional spherically averaged independent-atom models.

Peptide-bond anisotropy was analysed using the 0.97 Å dataset for PDB entry 6MU9, a high-resolution *β*-lactamase structure previously examined as a benchmark for peptide-bond chemistry (Panjikar & Weiss, 2025).

Within this structure, the Lys299–Glu300 peptide bond was selected as a positive-control site. Prior difference-density analysis identified the Lys299 carbonyl oxygen (O299) as best described by an O-protonated model, providing an experimentally motivated test of whether directional dipolar fingerprints are consistent with protonation-associated polarisation.

#### Backbone-wide dipolar anisotropy

Dipolar electron-density maps calculated at *ℓ* = 1 revealed reproducible, bond-centred anisotropy across the protein backbone. The paired lobes that straddle the peptide unit are aligned approximately with the local C–O and N–H bond directions, indicating that the angular decomposition captures the intrinsic polarity of the peptide linkage. This behaviour is consistent with the expected directional charge redistribution associated with carbonyl polarisation and amide donation.

#### Localisation at the Lys299 peptide bond

At the validated Lys299 site, the *ℓ* = 1 signal co-localised with the conventional *mF_o_*–*DF_c_* residual feature proximal to the carbonyl oxygen O299 (Fig. 7A,C). The dipolar lobes were concentrated on the O299–C299–N300 unit (i.e. the Lys299 carbonyl and the Glu300 amide N) (Fig. S14), supporting a localised electronic perturbation rather than a diffuse backbone feature.

**Figure 7:**
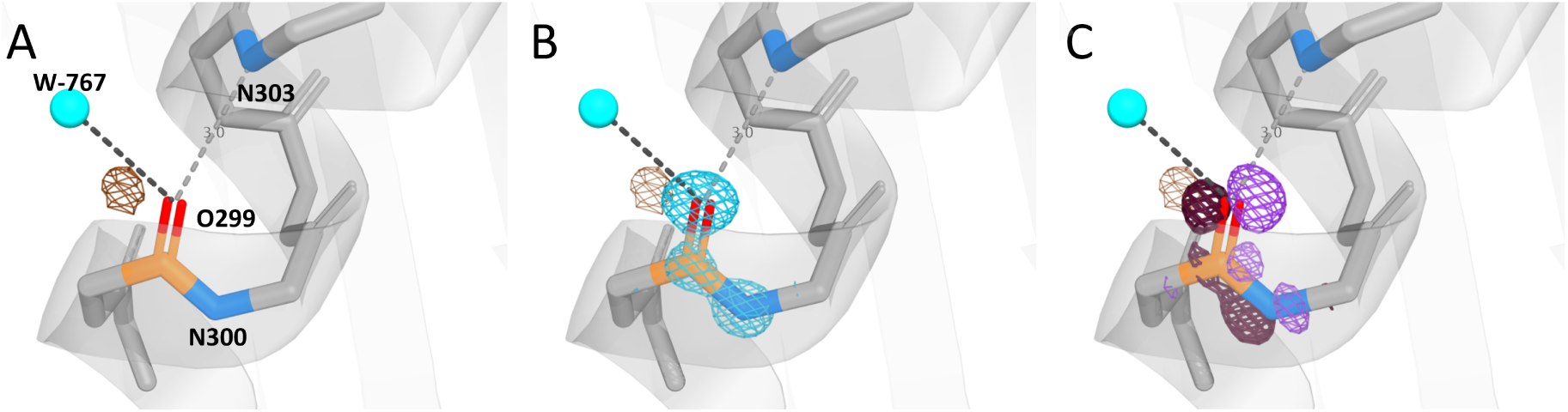
Angular decomposition of electron density at the 299–300 peptide bond in 6MU9. (A) Conventional difference map (*mF_o_* − *DF_c_*; mesh) around the peptide-bond region, showing residual density near the carbonyl oxygen O299 and neighbouring atoms including water W–767 and backbone N303. (B) Isotropic component (*ℓ* = 0), showing the approximately spherical density associated with the O299–C299–N300 peptide unit. (C) Dipolar component (*ℓ* = 1), displayed as paired positive and negative isosurfaces localised near the peptide bond. Positive and negative isosurfaces are contoured at the same absolute level. Contour levels were set to 2.8× the map r.m.s. For 6MU9, this corresponds to 0.392 e Å^−3^ for the *mF_o_* − *DF_c_* map (A) and 0.007 e Å^−3^ for the isotropic and dipolar component maps (B,C).

This localisation is consistent with the expected electrostatic response of a protonated or strongly polarised carbonyl environment, although the protonation state itself is inferred from complementary structural evidence. A stabilising hydrogen-bond environment was observed in the local structural context (Fig. 7A), including an O–H· · · O_water_ hydrogen bond to W–767 (∠O–H· · · O = 139.5^◦^, O· · · O = 2.79 Å) and a backbone interaction with N303. The residual peak adjacent to O299 is consistent with the O-protonated carbonyl assignment reported previously for this site (Panjikar & Weiss, 2025).

#### Directional structure of the dipolar component

The dipolar (*ℓ* = 1) component shows paired positive and negative lobes localised near the peptide bond (Fig. 7C). These features are oriented relative to the surrounding hydrogen-bond network, consistent with directional anisotropy in the local electron-density distribution. Figure 7 illustrates the separation of isotropic and directional contributions in a macromolecular environment. The *ℓ* = 0 component (Fig. 7B) reproduces the spherically averaged density envelope, whereas the *ℓ* = 1 component (Fig. 7C) isolates the leading directional anisotropy associated with the peptide unit.

#### Cartesian decomposition of the dipolar field

Decomposition into the three Cartesian directional components of the *ℓ* = 1 field is shown separately in Supplementary Fig. S14. These components correspond to (*p_x_, p_y_, p_z_*)-like directional modes aligned with the Cartesian axes, obtained from linear combinations of the complex spherical-harmonic functions *Y*_1,−1_, *Y*_1,0_ and *Y*_1, 1_ (Eqs. 6–8). Together, these directional components combine to form the rotationally interpretable dipolar vector field shown in Fig. 7C. The consistent localisation of signal at chemically meaningful positions supports both the numerical stability and the physical interpretability of the angular decomposition.

Individual *Y*_1*m*_ components depend on the choice of coordinate system and are therefore presented in the Supplementary material, while the combined *ℓ* = 1 dipolar field provides the physically interpretable directional representation.

#### Resolution and sampling dependence

To evaluate the robustness of these directional descriptors, resolution- and sampling-dependent behaviour was analysed systematically.

The stability of the inferred dipole axis depended on both the resolution cutoff (*d*_min_) and the local sampling radius (*r*_0_) used in the site-centred dipole estimation. For Lys299, the angle *θ* between the dipole axis and the local C–O bond varies across the *d*_min_ sweep, with the strongest variations occurring at the ends of the resolution range (Fig. 8A), and with a measurable dependence on the chosen sampling radius *r*_0_.

**Figure 8:**
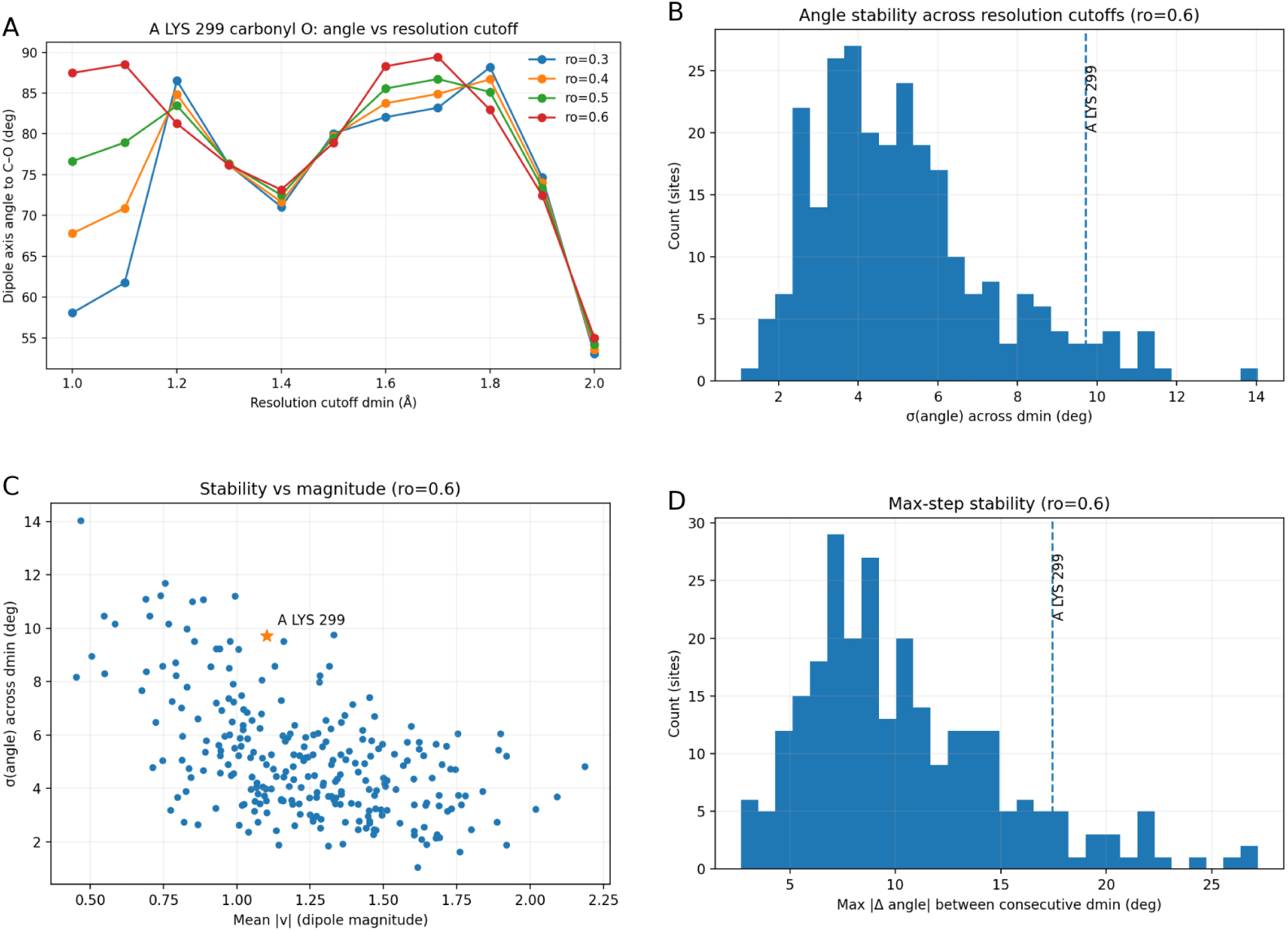
Resolution- and sampling-radius dependence of carbonyl-O dipole-axis orientation in 6MU9. **(A)** The dipole-axis angle *θ* of the Lys299 carbonyl oxygen relative to the local C–O bond is shown as a function of the resolution cutoff *d*_min_ for multiple local sampling radii *r*_0_ used in the dipole estimation. **(B)** Distribution of site-wise angular variability *σ*(*θ*) across the *d*_min_ sweep for all peptide carbonyl oxygens (shown for *r*_0_ = 0.6 Å), with Lys299 indicated (dashed line). **(C)** Relationship between *σ*(*θ*) and the mean dipole magnitude ⟨|*v*|⟩ across the *d*_min_ sweep (shown for *r*_0_ = 0.6 A), with Lys299 highlighted. **(D)** Distribution of the maximum stepwise angular change max |Δ*θ*| between consecutive *d*_min_ values (shown for *r*_0_ = 0.6 Å), with Lys299 indicated (dashed line). In (A), *d*_min_ increases left to right (i.e. progressively lower resolution); in (B,D), counts refer to the number of peptide carbonyl-O sites.

Using *r*_0_ = 0.6 Å, Lys299 lies in the high-variance tail of the site-wise distribution, with *σ*(*θ*) ≈ 10^◦^ across the *d*_min_ sweep (Fig. 8B), and a correspondingly elevated maximum stepwise change between successive cutoffs (max |Δ*θ*| ≈ 18^◦^; Fig. 8D).

#### Site-specific variability of dipole orientation

In the *σ*(*θ*)–⟨|*v*|⟩ relation (Fig. 8C), Lys299 combines substantial dipole magnitude with enhanced resolution dependence. This behaviour is consistent with an atypical local electrostatic environment or protonation-associated perturbation, rather than representing the backbone baseline.

#### Conservative assignment of light-atom features

Sites showing positive difference density near carbonyl oxygen were interpreted conservatively as candidate light-atom features. Assignment as an O-bound hydrogen was treated as provisional and considered reliable only when supported by refinement tests and compatible hydrogen-bond geometry, consistent with established crystallographic practice (Panjikar & Weiss, 2025).

The persistence of the dipolar signal in regions with moderate atomic displacement parameters indicates that the observed anisotropy is not an artefact of local ordering, atomic displacement parameters, or display choices. Instead, it reflects a robust component of the experimental electron density that can be partitioned into local chemical contributions and longer-range electrostatic coupling.

### 5.3 Aromatic residues and *π*-associated anisotropy

#### Aromatic residues as a test of delocalised anisotropy

Aromatic side chains provide chemical environments in which anisotropy may arise from delocalised *π*-electron structure rather than from simple bond polarity. These systems therefore provide a complementary test of whether the angular decomposition detects anisotropy beyond localised polar bonds.

#### Representative aromatic residues

To assess whether the angular decomposition captures *π*-associated anisotropy in a macromolecular context, dipolar (*ℓ* = 1) maps were examined for representative phenylalanine, tyrosine and tryptophan residues in high-resolution protein structures (Fig. 9), using examples Phe*85*, Tyr*210* and Trp*227* (chain A of 6MU9).

**Figure 9:**
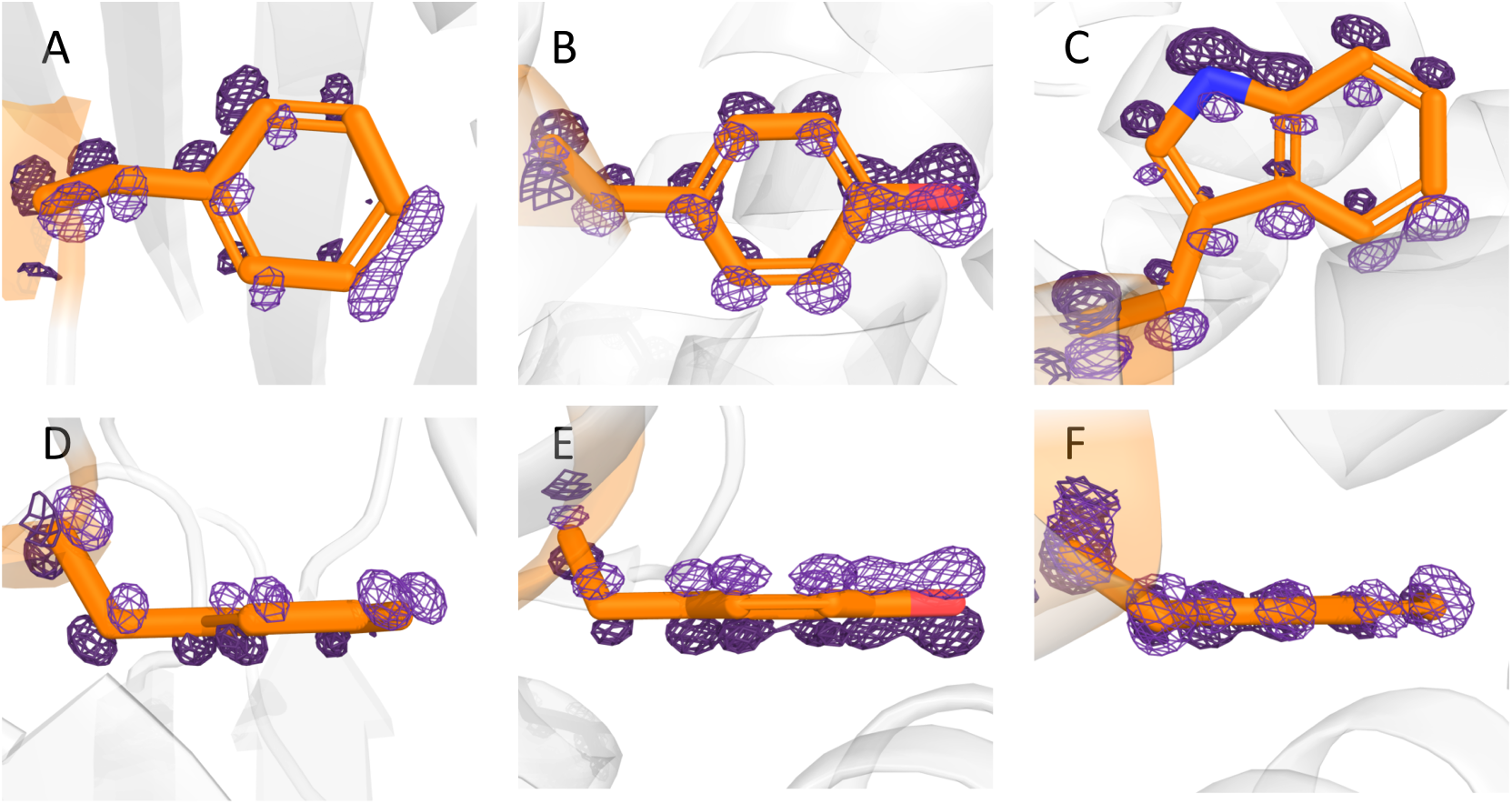
**Dipolar (***ℓ* = 1**) electron-density components associated with aromatic side chains.** (A–C) Face-on views of representative phenylalanine (A, Phe*85*), tyrosine (B, Tyr*210*) and tryptophan (C, Trp*227*) residues in 6MU9, viewed approximately along the aromatic ring normal. (D–F) Corresponding edge-on views for the same residues. Isomesh contours are drawn at symmetric positive and negative levels. For phenylalanine and tyrosine, maps are contoured at ±0.010 e Å, whereas for tryptophan the contour level is ±0.007 e Å . Positive and negative lobes are contoured at the same absolute level for each residue.

#### Observed dipolar features around aromatic rings

In all three residues, the *ℓ* = 1 maps display paired positive and negative lobes distributed around the aromatic ring. In tyrosine, lobes are observed above and below the ring plane, producing an approximately ring-normal pattern. In phenylalanine, the lobes are displaced both out of the ring plane and along the ring, indicating an oblique orientation relative to the ring normal. In tryptophan, the dominant *ℓ* = 1 density extends primarily within the ring plane and follows the extended conjugated aromatic system. These patterns are reproducible across chemically equivalent residues, indicating stable residue-specific anisotropy associated with aromatic environments.

#### Interpretation of aromatic anisotropy

The variation in dipolar orientation across aromatic residues indicates that the angular decomposition is sensitive not only to intrinsic aromatic *π*-electron structure, but also to modulation by local interactions such as hydrogen bonding, *π*–*π* stacking, and electrostatic contacts. Residues engaged in hydrogen bonding or intermolecular contacts display enhanced or distorted dipolar patterns, whereas solvent-exposed or weakly interacting rings show more symmetric, nearly ring-normal anisotropy.

#### Relative magnitude of aromatic dipolar signals

The magnitude of the aromatic dipolar signal is comparable to that observed for peptide N–H bonds, indicating that *π*-electron delocalisation produces directional anisotropy of similar strength to that generated by polar covalent bonds. Although no attempt is made here to extract quantitative ring-current or *π*-bond descriptors, these observations indicate that the angular decomposition is sensitive to both intrinsic *π*-electronic structure and its modulation by the surrounding protein matrix.

#### Integration with peptide-bond and benchmark observations

Taken together with the small-molecule benchmarks and peptide-bond analysis, the aromatic examples in Fig. 9 demonstrate that the *ℓ* = 1 component captures chemically distinct sources of anisotropy, including both localized polar bonds and delocalised *π* systems, within complex protein environments.

## 6 Discussion

### Scope and limitations

The present work focuses on the lowest-order aspherical contribution to the electron density (*ℓ* = 1). At typical macromolecular resolutions, dipolar anisotropy represents the dominant and most robust deviation from spherical symmetry, whereas higher-order terms (*ℓ* ≥ 2) are expected to be increasingly sensitive to noise, incomplete sampling and model phase uncertainty. The angular decomposition is therefore not intended as a replacement for multipolar refinement or quantum-crystallographic modelling, but as a complementary, model-independent analysis of directional electron-density features that can be extracted reliably from conventional diffraction data.

The method does not introduce additional refinement parameters and does not modify the underlying crystallographic model.

Because the method operates on experimentally refined structure factor amplitudes and phases without introducing additional adjustable parameters, the extracted anisotropic features reflect information already present in the diffraction data rather than artefacts introduced by model fitting, as confirmed by the numerical validation tests described in Supplementary Section S1.

Resolution limits, symmetry consistency and Friedel pairing were explicitly examined to ensure that the observed dipolar features are physically meaningful within the information content of the data. These properties were verified through controlled synthetic tests demonstrating rotational invariance, translational stability and conservation of angular power (Supplementary Section S1, Figs. S4–S7).

At the same time, the angular decomposition is constrained by the quality and completeness of the experimental structure factor amplitudes from which the maps are derived. The magnitude of the dipolar (*ℓ* = 1) signal becomes progressively less stable with decreasing resolution, as illustrated by the resolution-dependent behaviour shown in Fig. 8 and in the synthetic validation tests (Supplementary Section S1).

While the present implementation is robust for well-ordered regions of high-resolution structures, the interpretation of angularly filtered features in highly flexible or poorly ordered regions remains challenging and should be treated with appropriate caution. Resolution-dependent behaviour of angular power components is further illustrated in the synthetic validation tests presented in Supplementary Section S1.

The present study is complementary to, but distinct from, the analysis reported in *Peptide bond revisited*, in which the same high-resolution diffraction data (PDB entry 6MU9) were examined to assess protonation and partial double-bond character of peptide bonds through conventional difference-density analysis. Here, no modification of the crystallographic model or refinement strategy is introduced; instead, directional information is extracted directly from the experimentally determined structure factor amplitudes by angular decomposition. Whereas the earlier work focused on identifying specific chemical states within a refined electron-density map, the present approach interrogates how anisotropic electronic features are encoded in reciprocal space prior to any atom-centred interpretation independently of any atom-centred model assumptions. The two analyses therefore address different physical questions using the same underlying data, and their combined use highlights the richness of information accessible from high-resolution macromolecular diffraction experiments without invoking parameter-heavy models or additional experimental input.

The consistency of dipolar features observed across datasets refined using different modelling strategies (HAR and IAM) further indicates that the angular decomposition probes directional information intrinsic to the experimental structure factors rather than features imposed by a specific refinement model.

For aromatic side chains, the dominant *ℓ* = 1 axis itself becomes a chemically informative descriptor, varying systematically between ring-normal (tyrosine), oblique (phenylalanine) and largely in-plane (tryptophan) orientations, and thereby reporting how local *π*-electron delocalisation and bond polarity are modulated by the surrounding environment.

Importantly, the present framework provides quantitative descriptors of directional structure through angular power spectra and component-specific amplitudes. These quantities can be compared across datasets, molecular environments, and experimental conditions. This enables anisotropic features to be analysed in a reproducible and statistically interpretable manner, rather than only as qualitative visualisations.

Together, these observations suggest a broader framework in which chemically meaningful electronic features can be accessed through complementary real- and reciprocal-space analyses of macromolecular diffraction data. In this context, angular decomposition provides a practical route for extracting directional electronic information from conventional crystallographic experiments without introducing additional model complexity.

These findings confirm that the spherical-harmonic formalism introduced in the Mathematical Framework can be applied directly to experimentally determined structure factors to extract chemically meaningful directional information.

The present framework therefore provides a practical experimental pathway to accessing directional electronic structure encoded in crystallographic structure-factor amplitudes, without reliance on multipolar refinement or quantum-chemical modelling. These conclusions are supported by the reproducible dipolar signatures observed in peptide bonds (Fig. 7) and aromatic residues (Fig. 9).

### Outlook

Future extensions will focus on systematic analysis of higher-order angular components (*ℓ* ≥ 2) and on the use of chemically defined local reference frames, enabling more detailed characterisation of anisotropic electron distributions when supported by data quality and spatial resolution. The incorporation of local coordinate systems anchored to bond directions and functional-group geometry will allow directional features to be interpreted relative to chemically meaningful axes, including ligand orientation, bond polarity, and functional-group symmetry.

In particular, higher-*ℓ* channels are expected to separate quadrupolar and higher-order features associated with *π*-bonding, lone-pair geometry, and multi-centre electron distributions from the dipolar polarisation captured at *ℓ* = 1. Such separation will increase chemical specificity in cases where the dipolar signal alone is ambiguous, and may enable direct identification of anisotropic features associated with metal coordination, conjugated systems, and hydrogen-bond networks.

A further important direction involves linking angular decomposition to electrostatic and protonation-dependent properties. Because dipolar and higher-order anisotropic components reflect directional charge redistribution, the present framework provides a natural basis for detecting protonation-state changes, hydrogen-bond polarity, and environment-dependent charge modulation. These capabilities may enable experimental electron-density data to be used directly to infer chemically relevant protonation and electrostatic features in macromolecular systems.

More broadly, angular decomposition of experimental structure-factor amplitudes provides a simple, model-independent route to visualising directional anisotropy in macromolecular electron-density maps. By linking conventional crystallographic Fourier synthesis to an angular-momentum description, the approach enables bond polarity and related anisotropic features to be analysed directly from diffraction data without the need for multipolar refinement.

In this framework, directional electron-density information becomes accessible as a quantitative observable, supporting the formulation of testable, chemistry-driven mechanistic hypotheses from high-resolution structures. The method therefore provides a practical bridge between conventional crystallographic map interpretation and chemically informed electronic-structure reasoning.

Potential applications include the identification of protonation-dependent polarisation effects in active sites, characterisation of hydrogen-bond directionality and strength, detection of environment-dependent modulation of aromatic and metal-associated electron distributions, integration with electrostatic-potential reconstruction and charge-density analysis, and the development of automated workflows for chemically guided interpretation of high-resolution structural data.

Future developments will also explore computational optimisation and scalable implementations suitable for large macromolecular datasets, supporting routine application of angular decomposition methods to modern structural biology workflows.

## Appendix A: Continuous spherical formulation of the angular decomposition

The angular decomposition developed in the main text is derived from the continuous representation of the electron density using the spherical Fourier–Bessel formalism. This appendix outlines the mathematical steps connecting the conventional plane-wave expansion to the discrete angular filtering expression used in Eq. (4).

### A1. Plane-wave expansion

Electron density may be expressed as a superposition of plane waves in reciprocal space,

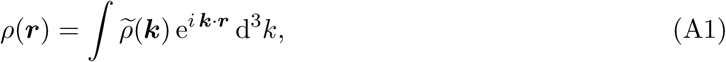

where *ρ*(***k***) denotes the continuous reciprocal-space representation of the density.

The plane-wave factor e*^i^* ***^k^***^·^***^r^*** may be expanded in spherical coordinates using the Rayleigh identity,

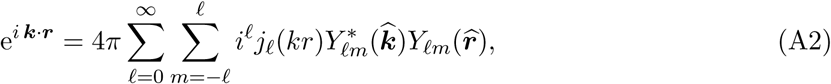

where *j_ℓ_*(*kr*) are spherical Bessel functions and *Y_ℓm_*are spherical harmonics.

Here ***k*** denotes a reciprocal-space wavevector expressed in Cartesian reciprocal coordinates. In crystallographic applications, allowed ***k*** values correspond to reciprocal-lattice vectors indexed by Miller indices.

### A2. Continuous angular decomposition

Substituting Eq. (A2) into Eq. (A1) yields

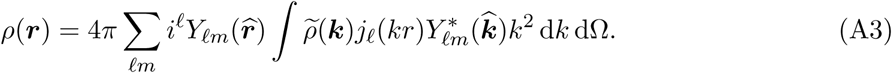

In the continuous spherical formulation, the functions *j_ℓ_*(*kr*) describe radial oscillations associated with each angular order. In the discrete crystallographic implementation adopted here, the radial dependence is retained implicitly through the conventional Fourier phase factors e^2^*^πi^ **^h^***^·***r***^. Consequently, explicit evaluation of spherical Bessel functions is not required, and angular filtering can be performed directly within the standard Fourier-synthesis framework.

This expression separates radial and angular dependence of the reciprocal-space density. The angular coefficients are therefore obtained through projection onto spherical harmonics,

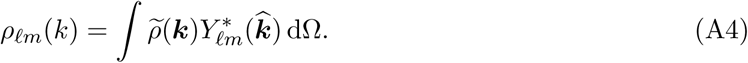

integration is over directions only. For each fixed reciprocal-space radius *k*, the angular coefficients are obtained by projection onto spherical harmonics over the angular variables only,

### A3. Radial shell representation

The integral over reciprocal space in Eq. (A3) contains the volume element

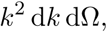

which corresponds to integration over spherical shells of radius *k*.

In the discrete crystallographic case, structure factors are available only at reciprocal-lattice points. The continuous radial integral is therefore approximated by partitioning reciprocal space into finite shells of thickness Δ*k*.

The contribution of shell *s* is approximated by

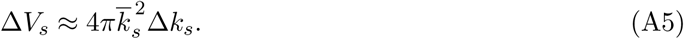

### A4. Discrete approximation for crystallographic data

In crystallographic Fourier synthesis, the reciprocal-space integral is replaced by a sum over discrete Miller indices ***h***.

Angular projection of structure factors therefore becomes

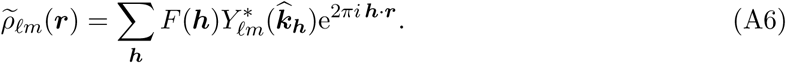

To approximate the continuous radial integration, each reflection is assigned a shell-dependent Weight

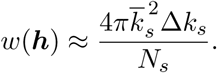

which corresponds to the shell-weight definition given in Eq. (5) of the main text.

### A5. Recovery of the discrete synthesis expression

Including the shell-weight normalization yields the final discrete form,

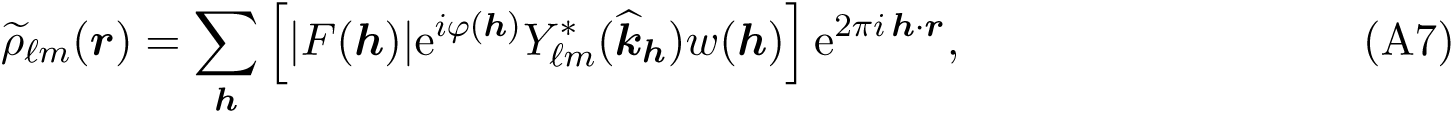

where ***r*** denotes fractional coordinates within the crystallographic unit cell, |*F* (***h***)| and *φ*(***h***) are the measured structure-factor amplitudes and phases, ***k_h_*** is the unit direction of the reciprocal-space vector associated with Miller indices ***h***, and *w*(***h***) is the shell-weight normalization defined in Eq. (5) of the main text.

Equation (A7) is identical to Eq. (4) of the main text. This result demonstrates that the angularly filtered Fourier synthesis used in the discrete crystallographic implementation constitutes a direct Riemann approximation to the continuous spherical Fourier–Bessel formulation derived above. This establishes that the discrete angular filtering procedure implemented through standard Fourier synthesis retains the mathematical structure of the continuous spherical formulation while remaining compatible with experimental crystallographic data.

## Supporting information

Synthetic validation tests and supplementary methods for the spherical-harmonic decomposition.

## Acknowledgements

Support from the Australian Synchrotron (ANSTO) is gratefully acknowledged. Computational resources used in this work were provided through institutional computing infrastructure. Helpful discussions with members of the crystallographic computing community are gratefully acknowledged. The author gratefully acknowledges Kay Diederichs for careful reading of the manuscript and for insightful comments that significantly improved the clarity of the presentation. The author also gratefully acknowledges P. Bhanu, Paul Tucker, Paulina Domniak, and Andy Karplus for helpful discussions at earlier stages regarding real-space approaches related to this work.

## Data availability

All datasets used in this study are publicly available. Small-molecule datasets were obtained from the Cambridge Structural Database (CSD): GLYALB05 (CCDC deposition number 995876) and UREA01. Macromolecular data were obtained from the Protein Data Bank (PDB), entry 6MU9.

