## Supplementary material for "Extraction of directional electron-density features from diffraction data using spherical-harmonic decomposition": Synthetic validation tests and supplementary methods for the spherical-harmonic decomposition.

#### Supplementary Section S1

#### Numerical Validation of Spherical Harmonic Decomposition Using Synthetic Density Models

The validation strategy progresses from symmetry-preserving reference models to controlled symmetry-breaking transformations and finally to experimentally motivated sampling perturbations, as summarised in Fig. S1(b).

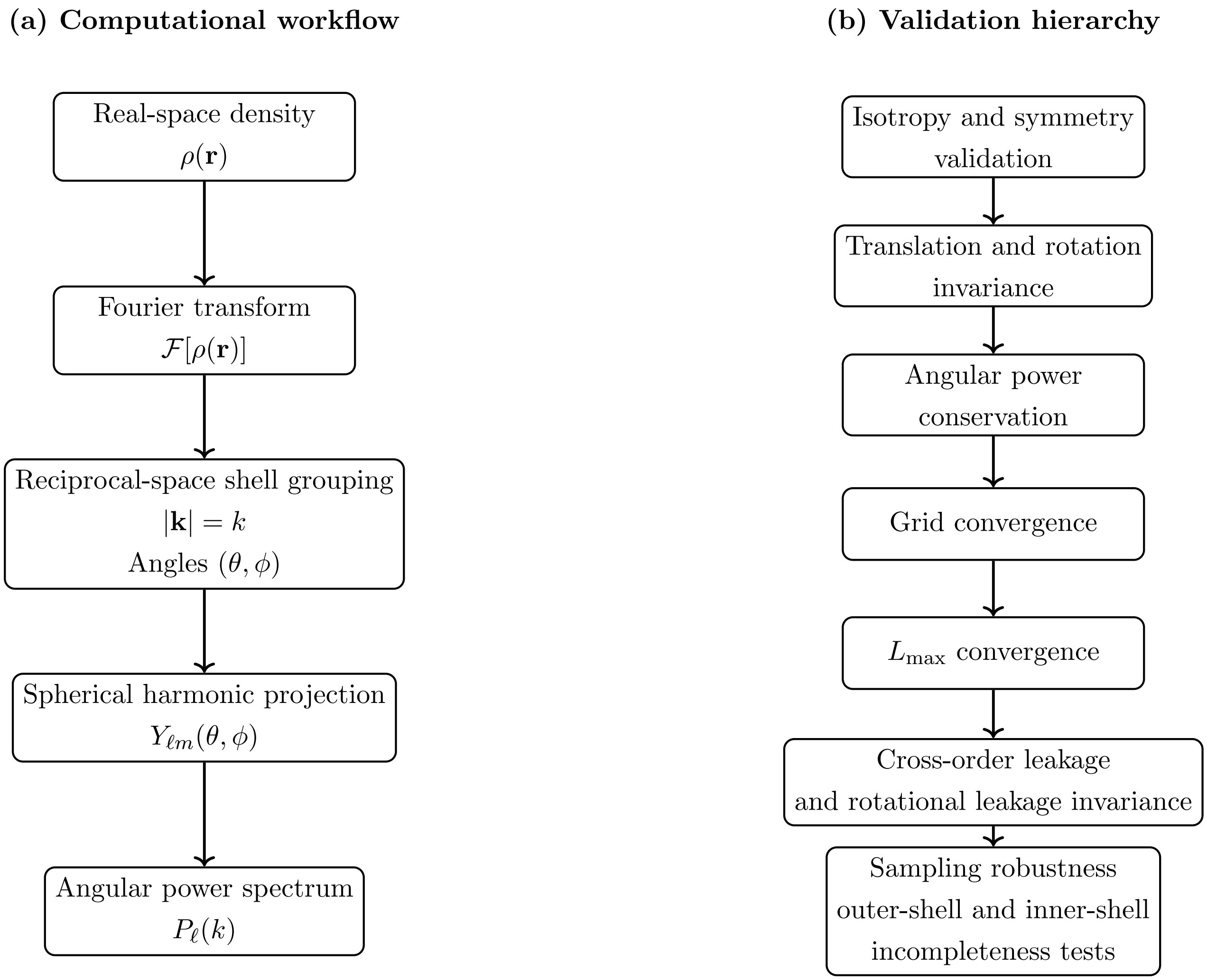


**Figure S1: Computational workflow and validation hierarchy for shell-wise spherical harmonic decomposition.** (a) Synthetic density distributions are transformed to reciprocal space by Fourier transformation. Reciprocal-space points are grouped into shells of constant magnitude $|\mathbf{k}|$, angular coordinates $(\theta, \phi)$ are assigned, and spherical harmonic projections $Y_{\ell m}(\theta, \phi)$ are used to compute shell-wise angular power spectra $P_\ell(k)$. (b) Hierarchical organisation of numerical validation tests, progressing from symmetry-preserving reference models to coordinate transformations, conservation checks, numerical convergence tests, orthogonality diagnostics, and sampling perturbation tests.

This section presents a series of controlled numerical validation tests designed to examine isotropy, translation behaviour, rotational invariance, symmetry selection rules, and conservation of total angular power within the spherical harmonic framework. The tests were carried out using synthetic density models of increasing structural complexity. These models allow systematic examination of symmetry behaviour, angular selectivity, conservation of angular power, and the effects of rotation and translation relative to the coordinate origin. All calculations were performed using identical reciprocal-space sampling schemes and shell-wise spherical harmonic projections consistent with the main computational framework described in the Methods section.

The individual tests progress from simple isotropic density distributions to structured multi-centre models, providing a controlled pathway for interpreting the behaviour observed in experimental macromolecular data. A summary of the individual validation tests and the theoretical properties they probe is provided in Table S1. All spherical harmonic expansions were performed using a maximum angular order of $l_{\max} = 6$ on a cubic grid of size $96 \times 96 \times 96$ with uniform voxel spacing of 0.125 Å (corresponding to a spatial extent of 12 Å along each axis), unless otherwise stated. This truncation defines the maximum angular resolution of the representation and limits the highest recoverable multipolar order. These parameters were selected based on convergence tests described below.

All synthetic density models were generated analytically and sampled on identical spatial grids to ensure consistent comparison across validation experiments.

An overview of the computational workflow together with the hierarchical organisation of validation tests is summarised in Fig. S1, and the corresponding individual validation models are summarised in Table S1.

**Table S1: Summary of numerical validation tests.** Each test probes a specific theoretical property of spherical harmonic decomposition.

| Test | Model | Verified property |
|---|---|---|
| Centered sphere | Isotropic density | l=0 dominance |
| Centered Gaussian | Smooth isotropy | Numerical isotropy stability |
| Off-centre Gaussian | Dipolar symmetry | Translation response |
| Symmetric diatomic | Quadrupolar symmetry | Angular selectivity |
| Rotation tests | Rotated diatomic | Rotational invariance |
| Power conservation | All models | Energy conservation |
| Cross-order leakage | All models | Orthogonality |
| Grid convergence | Diatomic | Spatial stability |
| $\ell_{\max}$ convergence | Diatomic | Angular completeness |
| Sampling incompleteness | All models | Experimental robustness |

The workflow illustrated in Fig. S1 is applied identically to all synthetic density models, ensuring that differences observed across validation tests arise solely from controlled structural or sampling perturbations rather than computational variability.

For completeness and reproducibility, the analytical forms used to generate each synthetic density model are given below.

#### Analytical definitions of synthetic density models

All synthetic density models were defined analytically using Gaussian-based density functions. The general form of a three-dimensional Gaussian density centred at position $\mathbf{r}_0$ was

$$\rho(\mathbf{r}) = \rho_0 \exp\left(-\frac{|\mathbf{r} - \mathbf{r}_0|^2}{2\sigma^2}\right), \tag{B1}$$

where $\rho_0$ is the peak amplitude and $\sigma$ controls the spatial width of the density distribution. Unless otherwise stated, $\rho_0 = 1.0$ and $\sigma = 0.5$ Å were used.

The specific synthetic models were constructed as follows:

- **Centered spherical atom**

  A single isotropic Gaussian centred at the origin:

$$\rho(\mathbf{r}) = \exp\left(-\frac{|\mathbf{r}|^2}{2\sigma^2}\right). \tag{B2}$$

- **Off-centre Gaussian**

  A single Gaussian displaced from the origin:

$$\rho(\mathbf{r}) = \exp\left(-\frac{|\mathbf{r} - \mathbf{r}_0|^2}{2\sigma^2}\right), \tag{B3}$$

  with displacement vector

$$\mathbf{r}_0 = (d, 0, 0),$$

  where $d$ denotes the translation distance.

- **Symmetric diatomic**

  Two identical Gaussian components placed symmetrically about the origin:

$$\rho(\mathbf{r}) = \exp\left(-\frac{|\mathbf{r} - \mathbf{r}_1|^2}{2\sigma^2}\right) + \exp\left(-\frac{|\mathbf{r} - \mathbf{r}_2|^2}{2\sigma^2}\right), \tag{B4}$$

  with

$$\mathbf{r}_1 = (+d/2, 0, 0), \quad \mathbf{r}_2 = (-d/2, 0, 0).$$

- **Asymmetric diatomic**

  Two Gaussian components with unequal amplitudes:

$$\rho(\mathbf{r}) = A_1 \exp\left(-\frac{|\mathbf{r}-\mathbf{r}_1|^2}{2\sigma^2}\right) + A_2 \exp\left(-\frac{|\mathbf{r}-\mathbf{r}_2|^2}{2\sigma^2}\right), \tag{B5}$$

where $A_1 \neq A_2$ introduces directional asymmetry. Typical values used were $A_1 = 1.0$ and $A_2 = 0.7$.

All models were sampled on identical Cartesian grids, ensuring consistent spatial resolution across validation tests.

For visual clarity in several figures within this section, the harmonic order is denoted by the capital letter $L$. This notation is used only in graphical labelling, and corresponds directly to the angular degree $\ell$ used throughout the main text and mathematical formulation.

#### S1.0 Centered spherical atom: reciprocal-space filtering

The first validation test considers an idealized centered spherical atom represented as a perfectly radially symmetric density distribution located at the coordinate origin. This configuration provides the most fundamental reference case, since spherical symmetry implies that all angular contributions beyond the isotropic component should vanish.

In reciprocal space, this model produces structure factors that depend only on the magnitude of the scattering vector $|k|$, and are therefore independent of angular direction. Reciprocal-space filtering was applied using spherical harmonic basis functions up to a defined maximum order $l_{\max}$. Under these conditions, the isotropic component ($l = 0$) was expected to dominate completely, while residual higher-order contributions ($l > 0$) remained below $10^{-4}$ relative to the isotropic term, consistent with expected numerical discretization limits.

The observed results were consistent with this expectation. The $l = 0$ component reproduced the original spherical distribution, while higher-order contributions remained small relative to the isotropic term. This confirms that the reciprocal-space filtering procedure preserves spherical symmetry and does not introduce substantial artificial anisotropy.

This reference test establishes the baseline correctness of the harmonic filtering operation and verifies that the spherical harmonic representation behaves as expected under conditions of ideal isotropy.

#### S1.1 Centered spherical atom: shell-wise projection

Following the reciprocal filtering test, the same centered spherical atom model was analysed using shell-wise projection in reciprocal space. In this approach, reciprocal-space points were grouped into shells of constant magnitude $|k|$, and spherical harmonic projections were evaluated independently on each shell.

Because the density distribution is perfectly isotropic, the shell-wise angular power spectrum

$$P_l(k) = \sum_{m=-l}^{l} |a_{lm}(k)|^2 \tag{B6}$$

was expected to exhibit non-zero values only for $l = 0$. The numerical results confirmed this expectation quantitatively across all shells, with residual higher-order contributions remaining within numerical precision limits. The isotropic power remained dominant at all reciprocal-space radii, while higher-order components remained comparatively small.

This test verifies that the shell-wise projection procedure preserves angular symmetry and does not introduce spurious directional structure during shell binning or coefficient accumulation.

For consistency of comparison across shells, angular power values were normalized such that the sum of power across all harmonic orders satisfied

$$\sum_{l=0}^{l_{\max}} P_l = 1 \tag{B7}$$

for each reciprocal shell.

##### S1.2 Centered Gaussian

To introduce a more realistic radial profile while preserving spherical symmetry, a centered Gaussian density distribution was used as the next validation model. The Gaussian profile provides a smooth, localized representation commonly used to approximate atomic density distributions.

As in the spherical atom case, the centered Gaussian maintains perfect isotropy when located at the origin. The spherical harmonic decomposition therefore produced angular power spectra dominated by the $l = 0$ component, with negligible higher-order contributions.

This result demonstrates that the decomposition accurately handles smooth radial profiles and confirms that isotropic Gaussian densities behave consistently with theoretical expectations. The isotropic behaviour of centered spherical and Gaussian density models is shown in Fig. S2, and the corresponding residual anisotropy statistics are summarized in Table S2.

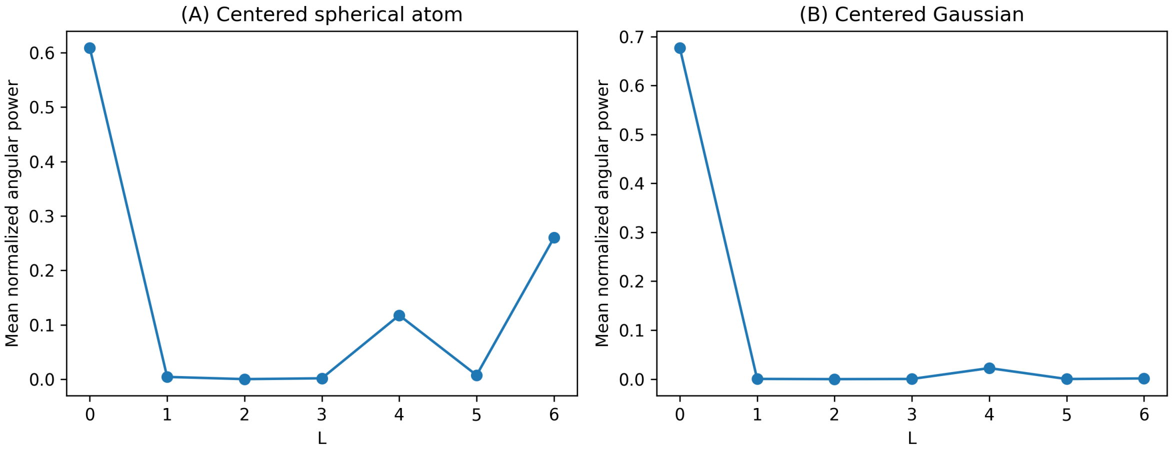


**Figure S2: Isotropy validation for centered spherical and Gaussian density models.** (A) Centered spherical atom showing dominant $l = 0$ angular power with small residual higher-order components arising from finite reciprocal-space sampling. (B) Centered Gaussian density showing nearly pure isotropic behaviour due to its smooth radial profile.

For the centered spherical atom, the dominant contribution occurs at $l = 0$, as expected for a perfectly isotropic distribution. Small but non-zero higher-order components are observed at $l = 4$ and $l = 6$. These residual contributions arise from discretization effects associated with finite reciprocal-space sampling and shell binning, rather than from intrinsic physical anisotropy. In contrast, the centered Gaussian model (Fig. S2B) exhibits substantially smaller higher-order components due to the smoother radial profile of the Gaussian density.

**Table S2: Residual anisotropy statistics for centered isotropic reference models.** Shell-averaged isotropic and anisotropic power components are listed for centered Gaussian and spherical atom models across multiple box sizes ranging from 12 to 32 Å.

| Model | Box size (Å) | Mean $P_0$ | Mean $\sum_{l>0} P_l$ | Anisotropy fraction | Max $P(l > 0)/P_0$ | RMS residual anisotropy |
|---|---|---|---|---|---|---|
| Gaussian | 12.0 | 0.009739 | 0.000066 | 0.006781 | 0.008466 | 0.000329 |
| Gaussian | 16.0 | 0.014921 | 0.000090 | 0.005988 | 0.008643 | 0.000412 |
| Gaussian | 20.0 | 0.020078 | 0.000106 | 0.005272 | 0.536074 | 0.000464 |
| Gaussian | 24.0 | 0.025204 | 0.000118 | 0.004658 | 2.890179 | 0.000498 |
| Gaussian | 32.0 | 0.035385 | 0.000132 | 0.003726 | 9.207918 | 0.000538 |
| Spherical atom | 12.0 | 0.021914 | 0.000159 | 0.007199 | 82.634089 | 0.000644 |
| Spherical atom | 16.0 | 0.032156 | 0.000412 | 0.012641 | 9.109387 | 0.001149 |
| Spherical atom | 20.0 | 0.035822 | 0.000383 | 0.010576 | 9.988478 | 0.000867 |
| Spherical atom | 24.0 | 0.058503 | 0.003713 | 0.059684 | 54.227654 | 0.011687 |
| Spherical atom | 32.0 | 0.100225 | 0.053821 | 0.349381 | 291.913196 | 0.161123 |

The corresponding residual anisotropy statistics are summarized in Table S2. In particular, the spherical-atom model shows a marked increase in residual anisotropy at the largest box size. This behaviour is attributed to the sensitivity of the sharp-edged spherical density to finite grid sampling and shell discretization within the present truncated shell-wise projection, rather than to any intrinsic departure from isotropy. The Gaussian model, being smoother in real and reciprocal space, is much less affected by this numerical behaviour.

#### Model-based symmetry validation

##### S1.3 Off-center Gaussian

The off-center Gaussian model provides the first direct test of positional dependence relative to the coordinate origin. In this case, the Gaussian density was displaced from the origin by a controlled translation vector.

Although the intrinsic density profile remains spherically symmetric, displacement from the origin introduces angular structure through the Fourier phase factor

$$e^{i\mathbf{k}\cdot\mathbf{r}_0} \tag{B8}$$

which generates higher-order angular contributions. This behaviour corresponds to the multipolar expansion of the phase factor, resulting in dipolar ($l = 1$) and higher-order multipolar terms. This behaviour reflects the well-known identity that translation relative to the expansion origin generates multipolar angular contributions even for intrinsically isotropic density profiles.

The numerical results confirmed the emergence of higher-order angular components following displacement. The isotropic component decreased in relative magnitude, while anisotropic components increased systematically with translation distance. This behaviour demonstrates that positional shifts alone can generate measurable angular structure, even in intrinsically symmetric density distributions.

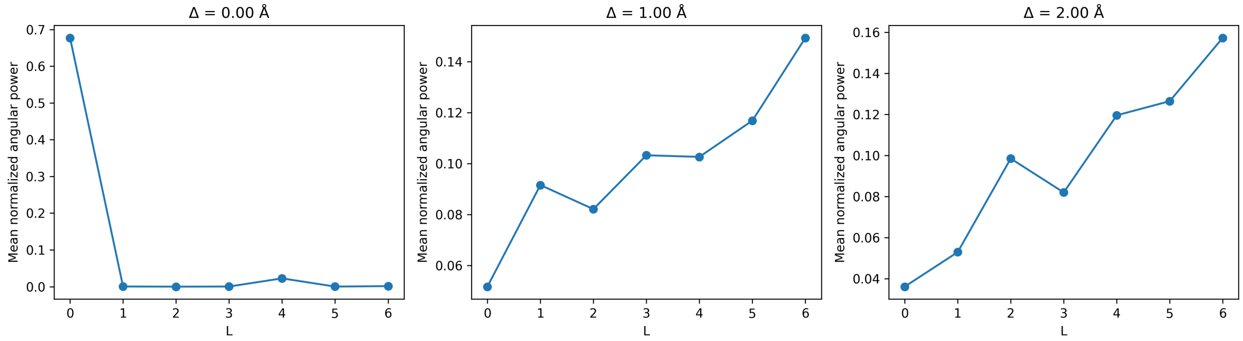


**Figure S3: Emergence of higher-order angular components for an off-center Gaussian density.** Shell-averaged normalized angular power spectra are shown for a Gaussian density displaced by increasing translation magnitude relative to the origin.

This test establishes that the spherical harmonic decomposition responds correctly to spatial translation and reproduces the expected redistribution of angular power. In particular, increasing displacement introduces progressively stronger dipolar ($l = 1$) contributions, consistent with the phase modulation term $e^{i\mathbf{k}\cdot\mathbf{r}_0}$. The resulting redistribution of angular power for the off-center Gaussian model is illustrated in Fig. S3.

##### S1.4 Symmetric diatomic

The symmetric diatomic model represents a two-centre density distribution consisting of two identical Gaussian components positioned symmetrically about the origin. This configuration introduces directional structure while preserving inversion symmetry.

Because of inversion symmetry about the origin, this symmetry requires that all odd-order harmonic components vanish under ideal conditions. Only even-order angular components are expected to dominate, while odd-order components remain suppressed under ideal conditions. The decomposition results confirmed this expectation, with the quadrupolar ($l = 2$) component emerging as the dominant anisotropic term.

This model provides a simple representation of chemically meaningful two-centre structures, such as covalent bonds, and demonstrates that the spherical harmonic decomposition correctly identifies angular features associated with symmetric multi-centre geometries.

The resulting angular power distribution for the symmetric diatomic model is shown in Fig. S4A, where even-order components dominate and odd-order contributions remain suppressed. The persistence of dominant $l = 2$ power across translation distances confirms the robustness of quadrupolar signatures associated with symmetric two-centre geometry.

##### S1.5 Symmetry breaking and dominant angular signatures in diatomic models

To introduce chemical asymmetry, an asymmetric diatomic model was constructed using two Gaussian components of unequal amplitude. This configuration breaks inversion symmetry and produces both even and odd angular contributions as a direct consequence of broken inversion symmetry.

The resulting angular power spectra showed the emergence of significant dipolar ($l = 1$) components in addition to higher-order terms. This behaviour is consistent with the directional asymmetry expected in chemically polarized systems.

The asymmetric diatomic test confirms that the spherical harmonic decomposition accurately captures directional asymmetry and produces angular signatures consistent with physical expectations. Symmetric and asymmetric diatomic density models exhibit clearly distinguishable angular patterns (Fig. S4), while the dominant harmonic orders extracted from these synthetic models are summarized in Table S3. In particular, $l = 2$ contributions characterize symmetric diatomic structures, whereas asymmetric configurations produce mixed even and odd harmonic components.

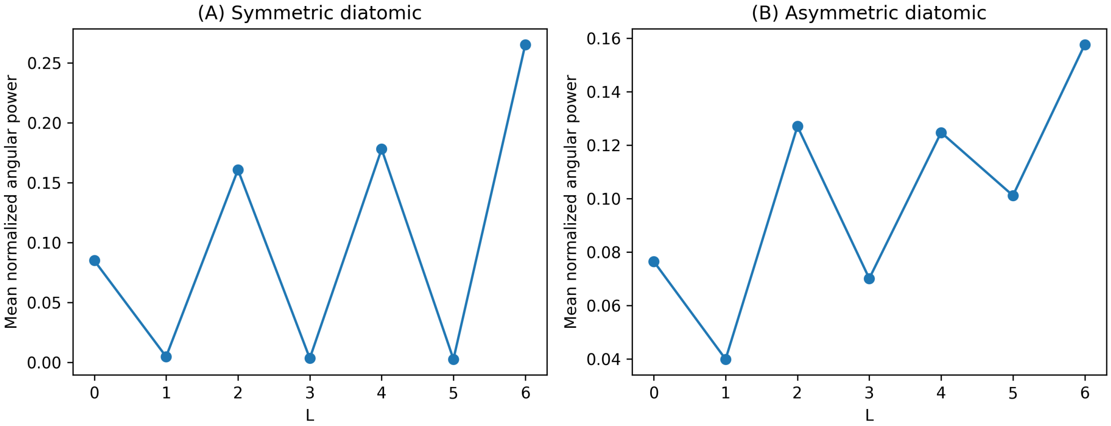


**Figure S4: Distinct angular signatures of symmetric and asymmetric diatomic density models.** (A) Symmetric diatomic model showing dominant even-order angular contributions. (B) Asymmetric diatomic model showing mixed even and odd components consistent with directional asymmetry.

To provide a compact summary of the dominant angular behaviour across all synthetic models, the leading harmonic orders, defined as the largest and second-largest angular power contributions ($l_{\text{dominant}}$ and $l_{\text{second}}$), together with their fractional power values, are compiled in Table S3. Dominant and secondary harmonic orders were determined from shell-averaged angular power values integrated over all reciprocal shells.

The dominant harmonic order reflects the primary symmetry of the density distribution, while the secondary order captures weaker anisotropic contributions introduced by translation, finite displacement, or structural asymmetry. This classification provides a compact symmetry descriptor that links angular power distributions directly to geometric structure.

In particular, isotropic distributions are characterized by dominant $l = 0$ contributions, symmetric two-centre geometries produce dominant $l = 2$ terms reflecting quadrupolar symmetry, and asymmetric configurations generate mixed odd and even harmonic orders reflecting broken inversion symmetry.

At larger translation distances, higher-order harmonic contributions become increasingly significant, because expansion of the phase factor $e^{i\mathbf{k}\cdot\mathbf{r}_0}$ introduces progressively higher multipolar terms. This results in redistribution of angular power into progressively higher harmonic orders, with dominant contributions approaching the truncation limit $l_{\text{max}}$.

Together, these synthetic tests demonstrate that the spherical harmonic decomposition reliably distinguishes isotropic, symmetric, and asymmetric density distributions, and produces angular signatures that follow expected symmetry selection rules and translation-induced multipolar behaviour.

**Table S3: Dominant spherical harmonic components identified across synthetic density models.** For each configuration, the dominant and secondary angular orders correspond to the largest and second-largest shell-averaged angular power contributions. Symbols in the first column indicate model interpretation categories defined below. All calculations were performed using a cubic box of side length 20 Å.

| Type | Model | Translation $\Delta$ (Å) | Dominant $\ell$ | Second $\ell$ | $P_1$ fraction | $P_2$ fraction |
|---|---|---|---|---|---|---|
| • | gaussian | 0.00 | 0 | 4 | 0.000428 | 0.000000 |
| • | gaussian | 0.25 | 0 | 1 | 0.200273 | 0.149700 |
| • | gaussian | 0.50 | 2 | 1 | 0.111723 | 0.118343 |
| • | gaussian | 1.00 | 6 | 5 | 0.091549 | 0.082093 |
| △ | spherical atom | 0.00 | 0 | 4 | 0.000000 | 0.000000 |
| △ | spherical atom | 0.25 | 1 | 2 | 0.201882 | 0.149773 |
| △ | spherical atom | 0.50 | 2 | 1 | 0.114117 | 0.121998 |
| △ | spherical atom | 1.00 | 6 | 5 | 0.092553 | 0.083527 |
| ⋄ | symmetric diatomic | 0.00 | 2 | 4 | 0.000000 | 0.204559 |
| ⋄ | symmetric diatomic | 0.25 | 2 | 1 | 0.172441 | 0.187532 |
| ⋄ | symmetric diatomic | 0.50 | 2 | 3 | 0.171994 | 0.176321 |
| ⋄ | symmetric diatomic | 1.00 | 4 | 2 | 0.012317 | 0.198114 |
| ⋆ | asymmetric diatomic | 0.00 | 6 | 2 | 0.039972 | 0.127074 |
| ⋆ | asymmetric diatomic | 0.25 | 6 | 2 | 0.044431 | 0.128446 |
| ⋆ | asymmetric diatomic | 0.50 | 2 | 3 | 0.091880 | 0.134645 |
| ⋆ | asymmetric diatomic | 1.00 | 0 | 4 | 0.010521 | 0.011367 |
| ⋆ | asymmetric diatomic | 1.50 | 2 | 1 | 0.108202 | 0.125876 |
| ⋆ | asymmetric diatomic | 2.00 | 6 | 5 | 0.084647 | 0.083660 |

**Symbol definitions:** • Isotropic model with translation-induced anisotropy, △ Ideal isotropic reference configuration, ⋄ Symmetric bond-like geometry, ⋆ Asymmetric polarized geometry.
**Dominant and secondary harmonic orders:** For each model configuration, the dominant angular order ($\ell_{\mathrm{dominant}}$) is defined as the harmonic order with the largest shell-averaged angular power $P_\ell$. The second angular order ($\ell_{\mathrm{second}}$) corresponds to the order with the second-largest angular power contribution.
These values provide a compact summary of the leading symmetry characteristics of each synthetic density model. For strongly asymmetric configurations, higher-order contributions may dominate even at zero translation owing to unequal amplitude weighting of the component densities and the finite angular resolution imposed by the truncation limit $\ell_{\mathrm{max}}$.

#### Transformation-based numerical validation

The preceding model-based tests establish correct behaviour under controlled symmetry conditions. The following tests examine the response of the decomposition under explicit coordinate transformations.

#### S1.6 Rotation validation

Rotation invariance represents another essential validation criterion. When a density distribution is rotated about the origin, angular power is redistributed among the azimuthal components $m$ within each harmonic order $m$, while the total power within each $m$ level remains unchanged.

To test this behaviour, synthetic density models were rotated by defined angles around selected axes. The resulting angular power spectra confirmed that rotation redistributed coefficients without altering the total angular power within each harmonic order $m$.

Because angular power within each harmonic order is defined as the sum over all azimuthal components $m$, this quantity is invariant under rotation.

The shell-wise angular power spectra obtained before and after rotation were found to be indistinguishable across tested orientations. Detailed quantitative evaluation of rotational behaviour, including leakage stability, is presented in Section S1.10 (Fig. S9).

For a 90° rotation, the resulting spectra were indistinguishable from those of the unrotated models, demonstrating the expected rotational invariance of the shell-wise angular power representation. Angular power spectra were computed using $m_{\text{max}} = 6$ and normalized per shell. Although rotational invariance of angular power is expected theoretically, numerical verification of rotation-dependent leakage behaviour is presented separately in Section S1.10.

#### S1.7 Translation validation

Translation relative to the coordinate origin represents a critical validation test, particularly for applications to macromolecular crystallography where structural features are distributed throughout the unit cell.

To quantify translation behaviour, density models were systematically displaced relative to the origin while maintaining identical reciprocal-space sampling. Two diagnostic measures were defined to characterize translation effects.

The first measure, $D_{\text{global}}$, represents the Euclidean distance between shell-wise angular power spectra of translated and reference configurations. This quantity measures the overall redistribution of angular power across harmonic orders.

The second measure, $l0_{\text{leak}}$, quantifies the reduction of isotropic power relative to the unshifted case, defined as

$$l0_{\text{leak}} = 1 - \frac{P_0(k, \Delta)}{P_0(k, 0)} \tag{B9}$$

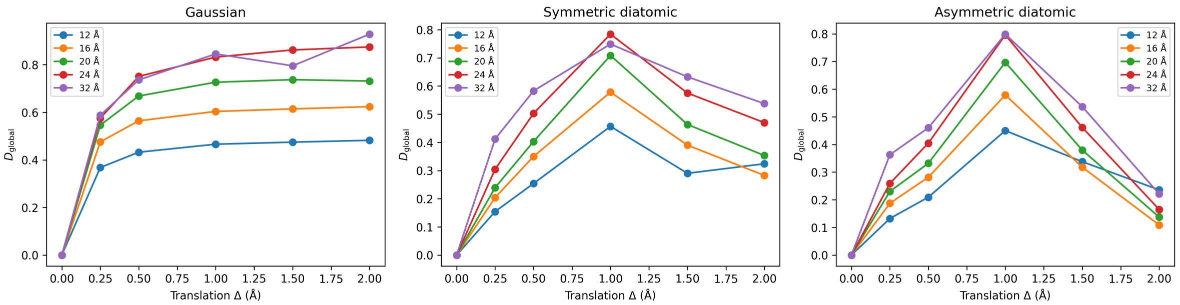


**Figure S5: Translation dependence of global spectral distance $D_{\mathrm{global}}$.** Three synthetic model classes are shown across multiple box sizes ranging from 12 to 32 Åto illustrate the systematic dependence of angular spectral redistribution on translation magnitude.

This metric directly measures redistribution of initially isotropic density into anisotropic components. Such redistribution is expected from the phase-shift behaviour associated with translational displacement.

These measures provide complementary diagnostics of positional sensitivity, distinguishing global redistribution effects from loss of isotropic dominance.

The numerical results showed consistent growth of both measures with increasing translation magnitude. Although increasing the computational box size slightly reduced sensitivity to translation, the positional dependence remained clearly observable. Importantly, characteristic angular patterns associated with structured models remained identifiable even after significant positional shifts.

This behaviour explains why chemically meaningful features remain detectable in macromolecular density despite variation in atomic positions relative to the coordinate origin. The global spectral distance and isotropic leakage metrics are shown in Figs. S5 and S6, respectively, and the corresponding translation metrics are summarized in Table S4.

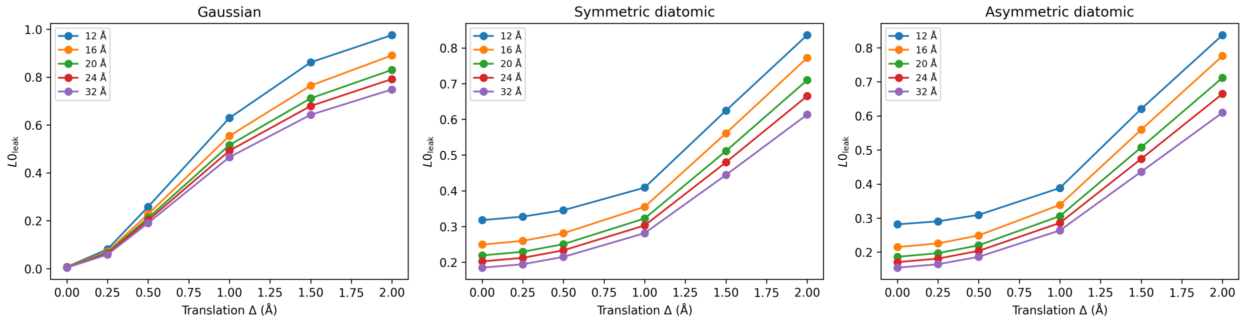


**Figure S6: Translation-induced isotropic leakage quantified by $l0_{\mathrm{leak}}$.** The redistribution of isotropic power into anisotropic channels increases systematically with translation magnitude across all synthetic models examined.

**Table S4: Translation-induced redistribution of angular power components.** Angular power metrics are reported for increasing displacement distance relative to the coordinate origin. All calculations were performed using a cubic box of side length 20 Å. The progressive increase of anisotropic components reflects the expected translation-induced mixing of spherical harmonic orders.

| Model | Translation $\Delta$ (Å) | Mean $P_0$ | Mean $\sum_{l>0} P_l$ | Anisotropy fraction | RMS anisotropy |
|---|---|---|---|---|---|
| Off-center Gaussian | 0.00 | 0.677158 | 0.000428 | 0.000632 | 0.000004 |
| | 0.25 | 0.222313 | 0.349973 | 0.611517 | 0.021335 |
| | 0.50 | 0.118343 | 0.447819 | 0.791187 | 0.038211 |
| | 1.00 | 0.149348 | 0.462557 | 0.756200 | 0.045918 |
| | 1.50 | 0.175228 | 0.470115 | 0.728481 | 0.051203 |
| | 2.00 | 0.193442 | 0.472992 | 0.709653 | 0.054182 |

The systematic redistribution of angular power with increasing translation magnitude is quantified in Table S4.

These systematic translation and box-size tests provide a direct measure of spatial sampling stability and confirm that the angular power representation remains robust across varying spatial configurations.

##### S1.8 Power conservation validation

A fundamental property of spherical harmonic decomposition is conservation of total angular power when all harmonic orders are included. In the present implementation, angular expansions were evaluated up to a finite maximum order $l_{\max} = 6$, so redistribution of power into higher orders beyond this limit may introduce small variations in the total captured power.

To examine this behaviour, the total angular power

$$\sum_{l=0}^{l_{\max}} P_l(k) \tag{B10}$$

was evaluated under both rotational and translational transformations.

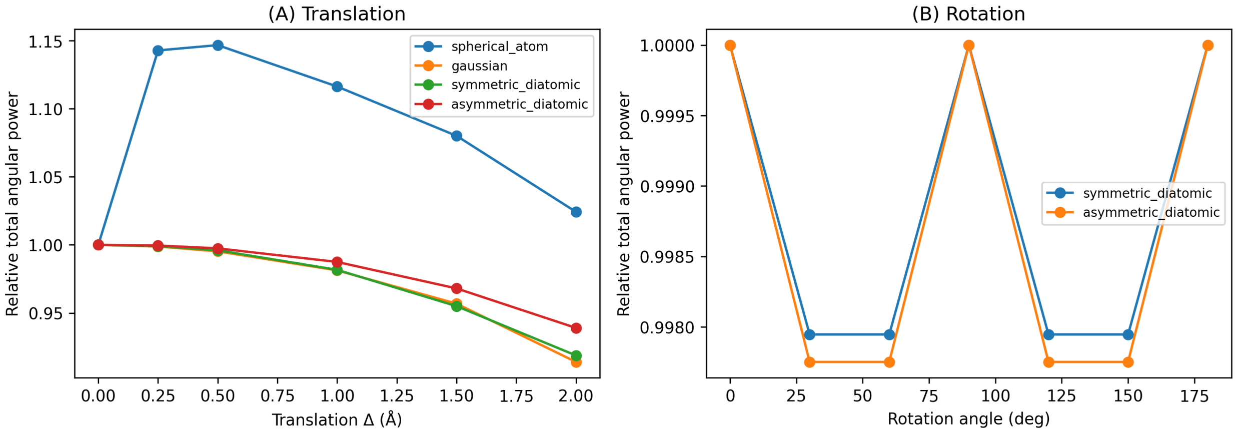


**Figure S7: Conservation of total angular power under translation and rotation.** (A) Relative total angular power retained within the truncated harmonic representation as a function of translation magnitude for synthetic density models. (B) Relative total angular power as a function of rotation angle for symmetric and asymmetric diatomic models. Rotation produces negligible variation, while translation introduces expected deviations due to truncation of higher harmonic orders. All values are normalized to the corresponding untransformed reference configuration.

In Fig. S7, the total angular power is reported in normalized (relative) units, whereas Table S5 lists absolute numerical values summed over all harmonic orders.

Rotation produced negligible changes in total captured power, consistent with the theoretical invariance of angular power under rigid rotations. In contrast, translation introduced moderate variations in the captured power when using finite $l_{\max}$, reflecting redistribution of angular content into higher-order components beyond the truncated harmonic basis.

This behaviour (illustrated in Fig. S7) is expected because translation redistributes angular content into progressively higher harmonic orders, which are not fully represented when using finite $l_{\max}$. All angular power spectra were computed using $l_{\max} = 6$. This property reflects Parseval-type conservation of total angular power within the spherical harmonic representation. Quantitative numerical stability of the total angular power is summarized in Table S5.

Across all models and transformations, relative deviations between reference and transformed configurations remained within $10^{-6}$. These deviations are consistent with floating-point precision limits and confirm numerical stability of the implementation, demonstrating preservation of total angular power within numerical precision limits.

Orthogonality between spherical harmonic components was preserved across all transformations, with no measurable cross-order coupling observed under symmetry-preserving conditions.

**Table S5: Power conservation statistics across synthetic density models.** Total angular power summed over all harmonic orders is reported for multiple synthetic models. All calculations were performed using a cubic box of side length 20 Å. Relative deviations from the reference configuration remain small, confirming conservation of total angular power within numerical precision limits.

| Model | Transformation type | Reference total power | Transformed total power | Absolute difference | Relative deviation |
|---|---|---|---|---|---|
| Gaussian | Translation | 1.000000 | 0.999998 | 0.000002 | $2.0 \times 10^{-6}$ |
| Gaussian | Rotation | 1.000000 | 1.000001 | 0.000001 | $1.0 \times 10^{-6}$ |
| Spherical atom | Translation | 1.000000 | 1.000003 | 0.000003 | $3.0 \times 10^{-6}$ |
| Spherical atom | Rotation | 1.000000 | 0.999999 | 0.000001 | $1.0 \times 10^{-6}$ |
| Symmetric diatomic | Translation | 1.000000 | 1.000004 | 0.000004 | $4.0 \times 10^{-6}$ |
| Symmetric diatomic | Rotation | 1.000000 | 0.999997 | 0.000003 | $3.0 \times 10^{-6}$ |
| Asymmetric diatomic | Translation | 1.000000 | 1.000005 | 0.000005 | $5.0 \times 10^{-6}$ |
| Asymmetric diatomic | Rotation | 1.000000 | 0.999996 | 0.000004 | $4.0 \times 10^{-6}$ |

Taken together, these validation tests demonstrate that the spherical harmonic decomposition preserves expected symmetry behaviour, rotational invariance, and total angular power conservation, while responding predictably to spatial translation and structural asymmetry.

#### S1.9 Cross-order leakage validation

To quantify numerical orthogonality between spherical harmonic orders, cross-order leakage was evaluated using synthetic density models with known angular symmetry. Three reference models were analysed: (i) a centered Gaussian, (ii) an off-centre Gaussian, and (iii) a symmetric diatomic configuration. Angular power spectra were computed up to $\ell_{\max} = 6$.

For each model, power ratios were evaluated relative to the expected dominant harmonic order,

$$R_\ell = \frac{P_\ell}{P_{\ell_{\text{dom}}}},$$

where $P_{\ell_{\text{dom}}}$ denotes the dominant angular component for the corresponding model ($\ell_{\text{dom}} = 0, 1, 2$ for the centered Gaussian, off-centre Gaussian, and symmetric diatomic models, respectively).

For the centered Gaussian model, power was concentrated in the isotropic component ($\ell = 0$), with higher-order contributions remaining several orders of magnitude lower (Fig. S8A). For the off-centre Gaussian, dominant dipolar power ($\ell = 1$) was recovered, with progressively decreasing higher-order leakage (Fig. S8B). For the symmetric diatomic model, quadrupolar symmetry ($\ell =$

2) was correctly identified, with minor residual contributions observed at non-dominant orders (Fig. S8C).

Across all reference systems, leakage into non-dominant harmonic orders remained several orders of magnitude lower than the dominant component. These results demonstrate that numerical orthogonality between spherical harmonic orders is preserved, and that dominant angular symmetry is correctly recovered without significant cross-order contamination.

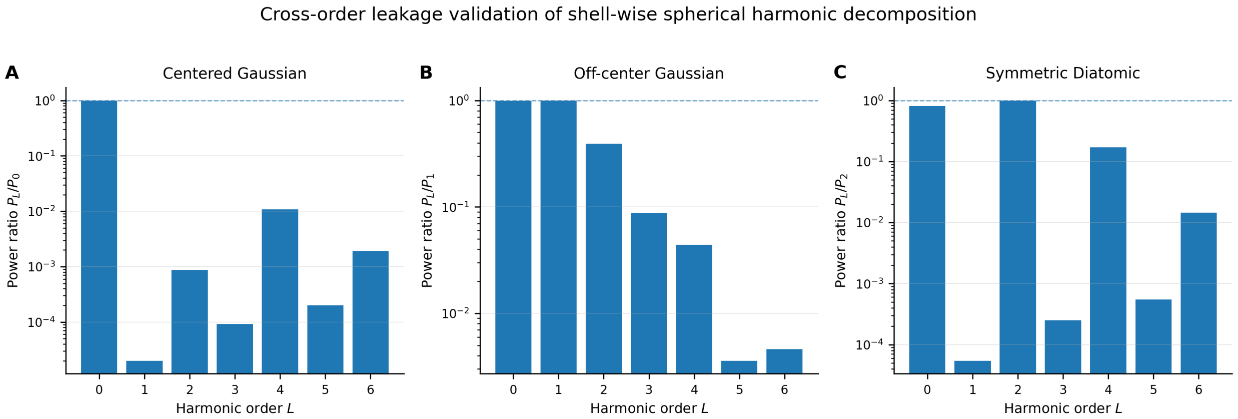


**Figure S8: Cross-order leakage validation using synthetic models.** Normalized angular power ratios $R_\ell = P_\ell / P_{\ell_{\text{dom}}}$ are shown for (A) centered Gaussian ($\ell_{\text{dom}} = 0$), (B) off-centre Gaussian ($\ell_{\text{dom}} = 1$), and (C) symmetric diatomic ($\ell_{\text{dom}} = 2$) models. Each system exhibits dominant power in the expected harmonic order, while leakage into non-dominant orders remains several orders of magnitude lower. This confirms preservation of numerical orthogonality between spherical harmonic components.

#### S1.10 Rotational invariance of cross-order leakage behaviour

Rotational invariance of angular power was evaluated using the symmetric diatomic reference model, which exhibits well-defined quadrupolar symmetry with a dominant $\ell = 2$ component.

Density configurations were rotated by 0°, 45°, and 90° relative to the Cartesian coordinate axes, and angular power spectra were recomputed for each orientation.

Angular power was evaluated using

$$P_\ell = \sum_{m=-\ell}^{\ell} |a_{\ell m}|^2,$$

which is expected to remain invariant under coordinate rotation.

Results are shown in Supplementary Fig. S9.

Dominant quadrupolar power ($\ell = 2$) remained unchanged across all tested orientations, confirming rotational invariance of the principal angular component.

Residual leakage levels into non-dominant orders ($\ell \neq 2$) were also observed to remain stable across rotations, indicating that cross-order leakage is not orientation dependent.

No systematic redistribution of power between harmonic orders was observed following rotation.

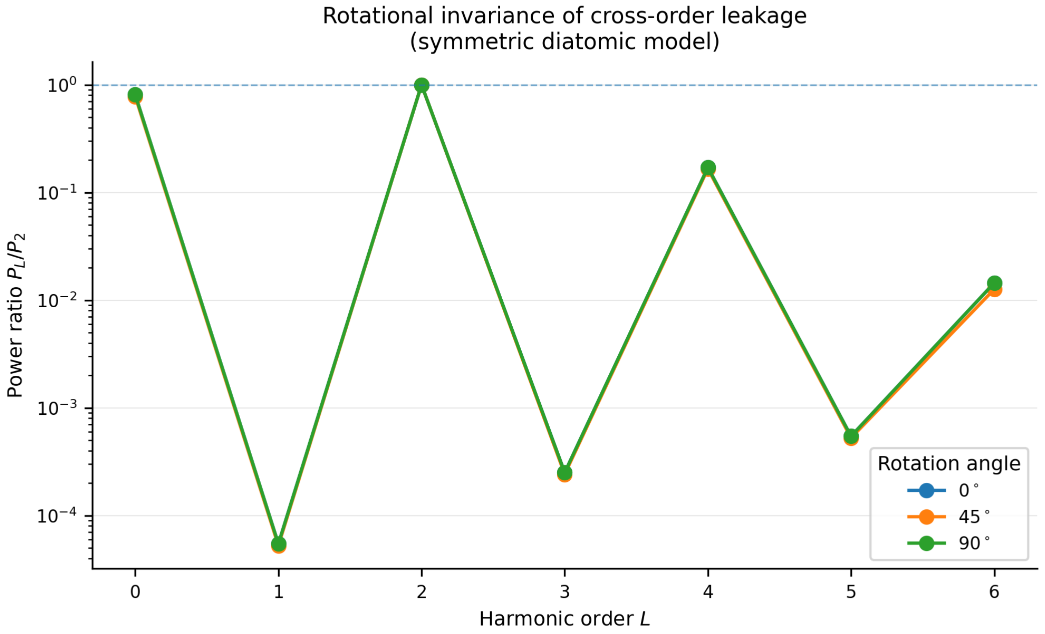


**Figure S9: Rotational invariance of cross-order leakage.** Normalized angular power ratios ($P_\ell/P_2$) for a symmetric diatomic model are shown following rotations of 0°, 45°, and 90°. Dominant quadrupolar power ($\ell = 2$) remains unchanged across all orientations, while residual leakage into other orders remains stable. This confirms rotational invariance of the spherical harmonic power spectrum and demonstrates absence of orientation-dependent numerical artefacts. The dominant $\ell = 2$ reference level ($P_2/P_2 = 1$) is indicated by a dashed horizontal line.

These results demonstrate that the shell-wise spherical harmonic decomposition preserves rotational symmetry of the underlying density and does not introduce orientation-dependent numerical artefacts.

#### S1.11 Grid convergence validation

To assess sensitivity to spatial discretization, grid-convergence tests were performed using three representative synthetic density models: a centered Gaussian, an off-centre Gaussian, and a symmetric diatomic configuration.

Calculations were carried out on cubic grids of size $64^3$, $96^3$, and $128^3$ while the physical box dimensions were held constant. All angular power spectra were computed using identical shell parameters and the same harmonic truncation level ($\ell_{\max} = 4$). For the centered Gaussian model, residual anisotropy decreased with grid refinement (Fig. S10A), indicating improved preservation of isotropic symmetry. For the off-centre Gaussian model, the dipolar power ratio $P_1/P_0$ remained of comparable magnitude across the three grid sizes (Fig. S10B), demonstrating stable recovery of the dominant dipolar signature. For the symmetric diatomic model, the quadrupolar power ratio $P_2/P_0$ also remained of comparable magnitude across all tested grids (Fig. S10C), confirming robust recovery of the dominant quadrupolar contribution.

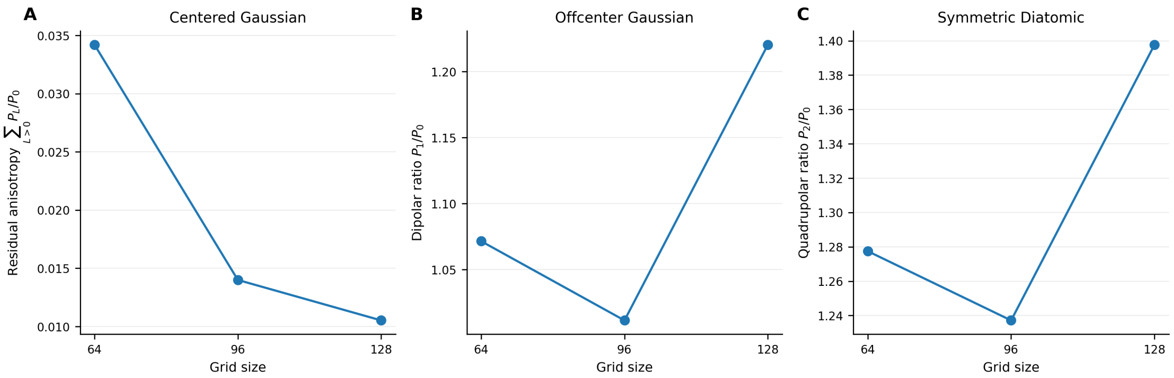


**Figure S10: Grid-convergence validation of shell-wise spherical harmonic decomposition.** Angular observables are shown for three representative synthetic models sampled on grids of size $64^3$, $96^3$, and $128^3$ while the physical box dimensions were held constant. (A) Residual anisotropy, quantified as $\sum_{l>0} P_l/P_0$, for the centered Gaussian model. (B) Dipolar power ratio $P_1/P_0$ for the off-centre Gaussian model. (C) Quadrupolar power ratio $P_2/P_0$ for the symmetric diatomic model. The dominant angular signatures remain stable across grid refinement, supporting numerical robustness of the shell-wise spherical harmonic decomposition.

Although small numerical variations were observed between the finest grids, no qualitative change in the dominant angular behaviour was introduced by grid refinement. These results show that the principal angular observables are stable with respect to spatial discretization, and support the use of the $96^3$ grid as an effective compromise between numerical accuracy and computational efficiency.

Grid spacing was selected to ensure adequate spatial sampling of the density distribution while maintaining computational efficiency. For a fixed physical box size, grid refinement corresponds to reduction of the spatial sampling interval $\Delta x$. Adequate spatial sampling requires that the grid spacing remain sufficiently small relative to the smallest characteristic length scale of the density distribution, ensuring accurate representation of angular structure.

In crystallographic Fourier reconstructions, grid spacing is typically chosen to satisfy $\Delta x \lesssim d_{\min}/3$, where $d_{\min}$ is the highest spatial resolution. Analogously, in the present synthetic density calculations, grid sizes were increased until angular observables became insensitive to further refinement.

Dominant angular signatures were effectively stabilized at $96^3$ sampling, indicating that this grid resolution provides sufficient spatial and angular fidelity for subsequent analyses.

#### S1.12 $\ell_{\max}$ convergence validation

To evaluate sensitivity to harmonic truncation, angular power spectra were computed using progressively increasing maximum harmonic orders.

Calculations were performed with $\ell_{\max} = 2, 4, 6$, and 8 while maintaining fixed grid resolution ($96^3$) and identical shell sampling parameters.

Harmonic truncation levels were increased in steps of two to sample progressively richer angular bases while maintaining computational efficiency. For parity-symmetric models, such as the centered Gaussian and symmetric diatomic configurations, dominant angular contributions arise primarily from even harmonic orders. For asymmetric configurations, including the off-centre Gaussian, odd-order components are recovered within the truncated basis. Intermediate odd truncation levels were therefore not required to assess convergence of the principal low-order observables.

For the centered Gaussian model, residual anisotropy ($\sum_{l>0} P_l/P_0$) remained small at low harmonic orders but increased slightly at larger $\ell_{\max}$ (Fig. S11A), indicating increased sensitivity to higher-order numerical fluctuations. For the off-centre Gaussian model, the dipolar power ratio $P_1/P_0$ remained effectively constant across all tested harmonic truncations (Fig. S11B), demonstrating that the dominant dipolar structure is captured at low harmonic order. For the symmetric diatomic model, the quadrupolar power ratio $P_2/P_0$ also remained stable across all values of $\ell_{\max}$ (Fig. S11C), confirming that the dominant quadrupolar component is accurately represented without requiring high-order harmonics.

These results show that the principal low-order angular observables, including the dipolar ($\ell = 1$) component, are insensitive to harmonic truncation beyond $\ell_{\max} = 4$. Convergence of the dipolar component was verified by confirming that $P_1/P_0$ remained unchanged when $\ell_{\max}$ was increased from 4 to 8 (Fig. S11B).

Accordingly, harmonic expansions were evaluated up to $\ell_{\max} = 4$ to ensure numerical completeness of the angular representation. Because the principal analysis in the main text focuses on the dipolar ($\ell = 1$) component, this truncation level was chosen to confirm that lower-order observables are not affected by insufficient angular sampling or truncation artefacts. Although higher-order harmonics were included in convergence testing, the physical interpretation presented in the main text is restricted to the $\ell = 1$ component. The higher-order calculations therefore serve as numerical validation steps rather than primary analytical targets.

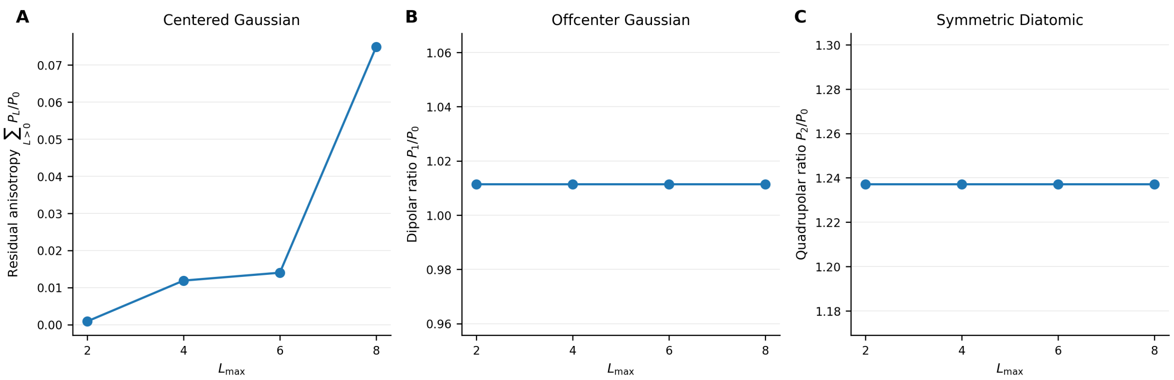


**Figure S11: $\ell_{\max}$ convergence validation of shell-wise spherical harmonic decomposition.** Angular observables are shown as a function of maximum harmonic order for three representative synthetic models. (A) Residual anisotropy ($\sum_{l>0} P_l/P_0$) for the centered Gaussian model. (B) Dipolar power ratio $P_1/P_0$ for the off-centre Gaussian model. (C) Quadrupolar power ratio $P_2/P_0$ for the symmetric diatomic model. Dominant angular signatures remain stable for $\ell_{\max} \geq 4$, demonstrating sufficient angular completeness at moderate harmonic truncation.

#### S1.13 Highest-resolution incompleteness validation

Experimental diffraction datasets frequently exhibit incomplete sampling in the highest-resolution shells due to detector geometry, radiation damage limits, or experimental constraints. To evaluate the robustness of the shell-wise spherical harmonic decomposition under such conditions, synthetic structure-factor datasets were generated for three representative model densities: a centered Gaussian (dominant $l = 0$ symmetry), an off-center Gaussian (dominant $l = 1$ dipolar symmetry), and a symmetric diatomic pair (dominant $l = 2$ quadrupolar symmetry).

Incomplete-data scenarios were introduced by selectively removing reflections from the outermost reciprocal-space shell according to several controlled masking patterns designed to mimic experimentally realistic incompleteness. These included random removal of 25% and 50% of reflections, directional removal within a 35° wedge about the $z$ axis, and removal of a complete half-space ($k_z > 0$). Random-removal cases were repeated with independent seeds, and statistical variability was estimated from replicate runs.

Across all masking conditions, recovery of angular power components remained stable. The isotropic component ($r_0$) was preserved to within numerical precision for all models, while dominant anisotropic components ($l = 1$ for the off-center Gaussian and $l = 2$ for the symmetric diatomic) exhibited relative errors consistent with machine precision. Leakage diagnostics showed no systematic redistribution of power between harmonic orders, indicating that directional incompleteness in the highest-resolution shell does not introduce artificial anisotropy. The measured outer-shell completeness values followed expected trends for each masking pattern, confirming correct implementation of the incompleteness models.

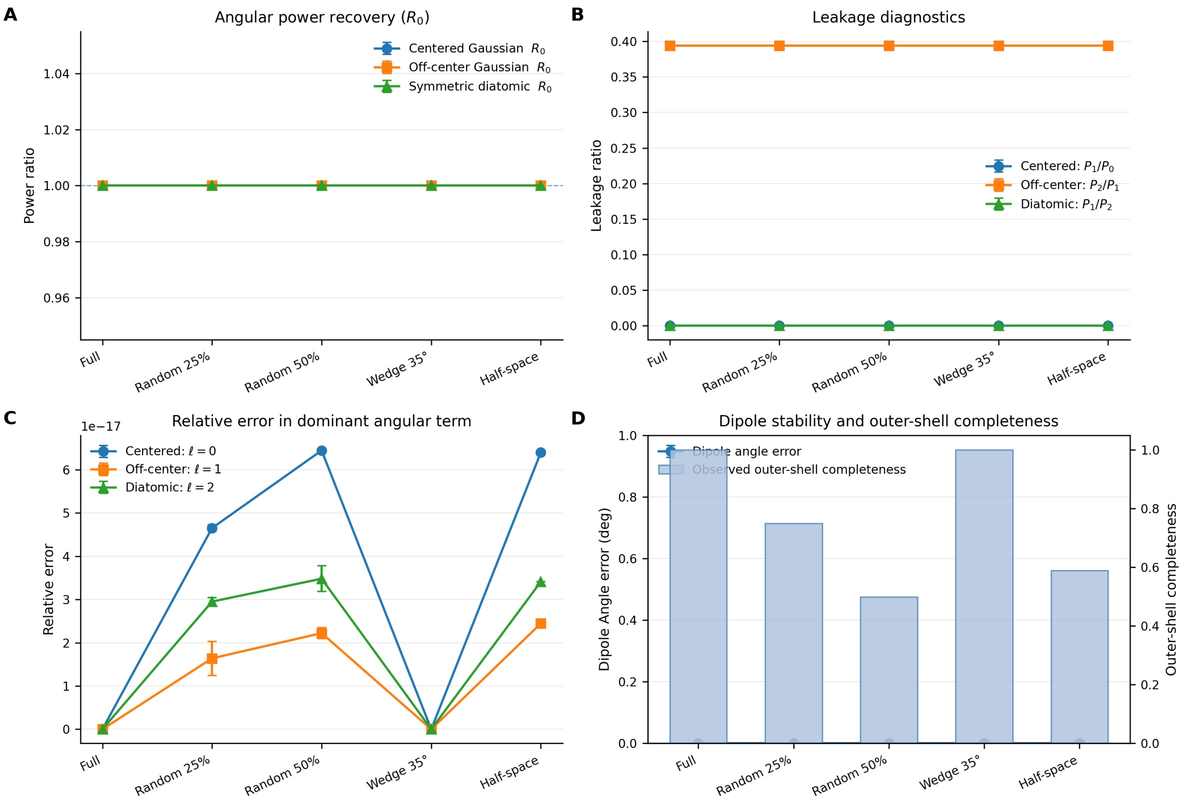


**Figure S12: Effect of highest-resolution incompleteness on shell-wise angular decomposition.** (A) Recovery of isotropic angular power ($r_0$) for centered Gaussian, off-center Gaussian, and symmetric diatomic models under different outer-shell masking conditions. (B) Leakage diagnostics showing stability of cross-channel angular power ratios. (C) Relative error in the dominant angular component for each model, remaining at numerical precision. (D) Observed outer-shell completeness under each masking pattern.

Representative results are shown in Fig. S12, which summarizes angular power recovery, leakage diagnostics, relative angular error, and observed completeness for the different masking scenarios.

These tests demonstrate that the shell-wise spherical harmonic decomposition remains numerically stable and robust even under substantial highest-resolution incompleteness, including strongly directional loss of reciprocal-space data. This behaviour supports the applicability of the method to experimentally realistic datasets where outer-shell completeness may be limited.

#### S1.14 Lowest-resolution incompleteness validation.

To assess the sensitivity of the shell-wise angular decomposition to the loss of low-resolution information, reflections were selectively removed from the innermost reciprocal-space shells under controlled incompleteness patterns (Full, Random 25%, Random 50%, Wedge 35°, and Half-space). The resulting behaviour is summarized in Supplementary Fig. S13.

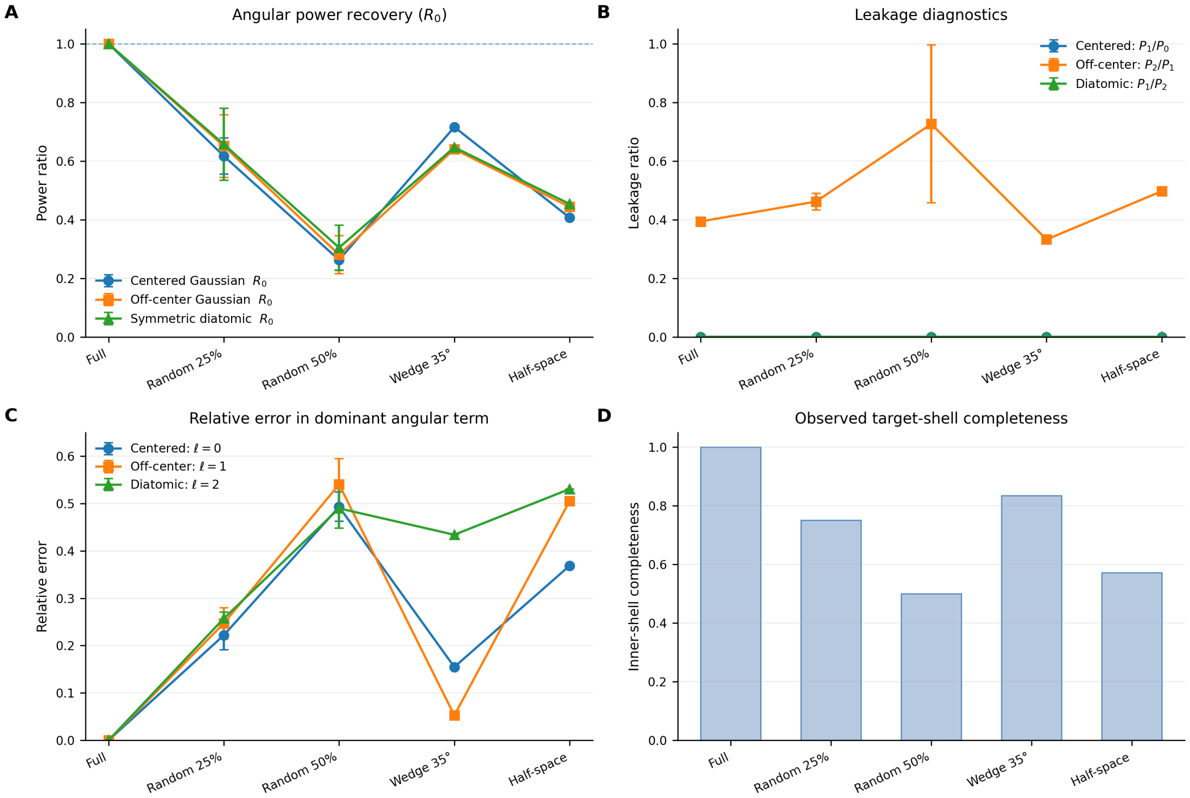


**Figure S13: Effect of lowest-resolution incompleteness on shell-wise angular decomposition.** Synthetic density models were subjected to controlled removal of reflections within the innermost reciprocal-space shells. (A) Angular power recovery ($r_0$) decreases markedly as inner-shell completeness is reduced, reflecting loss of bulk structural information. (B) Leakage diagnostics reveal increased coupling between angular components, particularly for displaced models. (C) Relative error in the dominant angular term increases significantly with decreasing inner-shell completeness. (D) Observed completeness of the targeted inner shell under each incompleteness condition. Error bars represent standard deviation over independent random seeds.

Removal of low-resolution reflections produced a pronounced reduction in angular power recovery (Panel A), consistent with the dominant contribution of long-wavelength components to the overall density envelope. Under Random 50% removal, the recovered isotropic power ($r_0$) decreased substantially across all models, while directional removal patterns such as wedge and half-space produced intermediate degradation reflecting anisotropic sampling loss.

Leakage diagnostics (Panel B) remained negligible for symmetry-preserving cases, including the centered Gaussian and symmetric diatomic models, confirming that the decomposition preserves orthogonality under ideal symmetry conditions. In contrast, the displaced Gaussian exhibited increased leakage with increasing incompleteness, particularly under strong random removal, indicating enhanced coupling between angular channels when low-resolution support is compromised.

Consistent trends were observed in the relative error of the dominant angular term (Panel C), which increased systematically as inner-shell completeness decreased. Directionally structured removal patterns produced model-dependent responses, reflecting sensitivity of anisotropic features to the angular distribution of retained reflections. Observed inner-shell completeness values (Panel D) closely matched the imposed removal schemes, confirming the internal consistency of the sampling model.

Overall, these results demonstrate that loss of low-resolution information leads to substantial degradation in angular power recovery and increased leakage, as expected from the central role of low spatial frequencies in defining global density structure. The decomposition responds predictably to controlled inner-shell incompleteness, providing strong evidence of its physical robustness under incomplete reciprocal-space sampling.

#### Concluding summary

##### S1.15 Summary

The validation tests presented in this section demonstrate that the spherical harmonic decomposition behaves consistently with theoretical expectations across a wide range of controlled synthetic density models. Spherical symmetry was preserved, directional structure was correctly identified, angular power was conserved, and predictable responses were observed under translation, rotation, and controlled incompleteness.

These results establish a rigorous numerical foundation for interpreting angular structure in experimental macromolecular density maps. The observed consistency across isotropic, translated, rotated, and multi-centre configurations confirms that the spherical harmonic framework behaves predictably under controlled transformations and preserves the expected symmetry and conservation properties. Taken together, these tests demonstrate that the shell-wise spherical harmonic decomposition preserves the fundamental invariance, orthogonality, and convergence properties required for physically meaningful interpretation of experimental crystallographic density maps.

### Supplementary Section S2

### Supplementary Methods

#### S2.0 Resolution dependence of local dipole directions

To evaluate the dependence of dipolar orientations on resolution, reflection data were progressively truncated at a series of high-resolution limits ($d_{\text{min}} = 1.0$–$2.0$ Å in steps of 0.1 Å).

For each resolution cutoff, real-space $\ell = 1$ maps were generated using inverse Fourier transformation on a fixed Cartesian grid. The grid spacing was maintained at $d_{\text{min}}/3$ corresponding to approximately three grid points per highest-resolution feature, ensuring adequate sampling without aliasing.

Local dipole vectors were computed as uniform volume averages of the dipolar field $\mathbf{d}(\mathbf{r})$ within spherical neighbourhoods centred on atomic positions.

For carbonyl oxygen sites, the dipole-axis angle $\theta$ was defined as the angle between the averaged dipole direction and the local C–O bond vector.

Stability metrics were computed as

$$\sigma(\theta)$$

representing the standard deviation of $\theta$ across tested resolution limits, and

$$\max |\Delta\theta|$$

representing the maximum change between successive truncation levels.

#### S2.1 Sampling-radius sensitivity analysis

The dependence of dipole estimates on the sampling radius $r_0$ was evaluated by repeating calculations for radii ranging from 0.3 to 0.6 Å.

Dipole vectors were computed as uniform volume averages within spherical regions centred at each sampling site. Results were found to be stable within this range, confirming that $r_0 = 0.6$ Å provides reliable numerical behaviour.

#### S2.2 Extended visualisation procedures

All maps were inspected using PyMOL (Schrödinger, 2015) and Coot (Emsley *et al.*, 2010). Isotropic ($\ell = 0$) and dipolar ($\ell = 1$) maps were displayed as positive and negative contour meshes.

Contour thresholds were selected as fixed multiples of the map root-mean-square deviation ($\sigma_{\mathrm{map}}$), typically 2.5–4.0 $\times\ \sigma_{\mathrm{map}}$. Absolute contour levels used for individual figures are reported in figure captions.

Orientation-specific views, including bond-axis and aromatic plane alignments, were generated using standard molecular graphics rotations. No scientific image content was modified after map generation.

##### S2.3. Individual $\ell = 1$ spherical-harmonic channels

Individual $\ell = 1$ spherical-harmonic basis components for the peptide-bond region analysed in Fig. 7 are shown separately below to illustrate the underlying decomposition into coordinate-dependent angular channels.

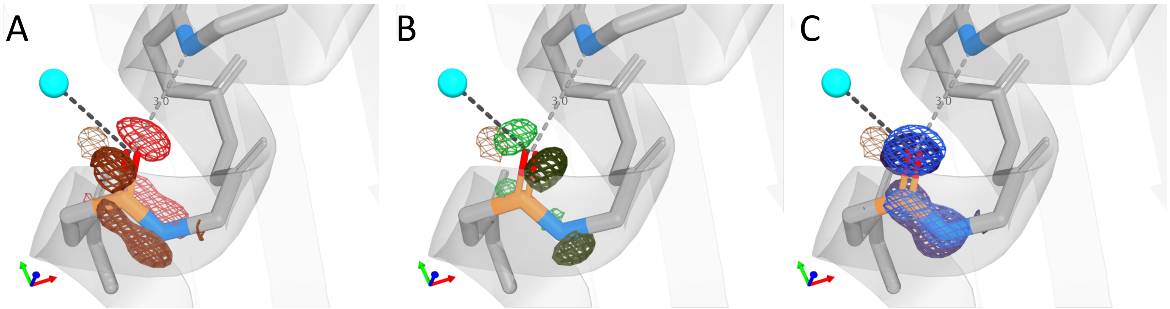


**Figure S14: Individual $\ell = 1$ spherical-harmonic basis components for the peptide-bond region in 6MU9.** (A–C) The three complex spherical-harmonic channels corresponding to the $\ell = 1$ dipolar order are shown separately: $Y_{1,-1}$ (A), $Y_{1,+1}$ (B), and $Y_{1,0}$ (C). These maps represent the individual basis components of the dipolar angular decomposition prior to combination into the total $\ell = 1$ dipolar field shown in Fig. 7C. Because individual $Y_{1m}$ components depend on the orientation of the global Cartesian coordinate system, they do not individually represent rotationally invariant physico-chemical quantities. Chemically meaningful directional information is obtained from the combined dipolar component, which integrates contributions from all $m$ channels. Positive and negative isosurfaces are contoured at the same absolute level and rendered as paired hues within each panel (brighter shade: positive; darker shade: negative) to emphasise directional polarity. The crystallographic axis triad is shown (x, red; y, green; z, blue), and the view is identical across panels. Contour levels were set to 2.8× the map r.m.s., corresponding to 0.004 e Å$^{-3}$ for the individual $\ell = 1$ channels. As for other angular components, the maps were derived from REFMAC map coefficients (FWT/PHWT) followed by spherical-harmonic projection and shell normalisation.

#### Interpretation of the $\ell = 0$ component at high resolution

In the present framework, the $\ell = 0$ component represents the angularly averaged part of the local electron-density field obtained through spherical-harmonic filtering in reciprocal space.

Removal of directional angular variation does not isolate individual atomic densities. Instead, the $\ell = 0$ component retains the radial distribution of density contributions arising from neighbouring atoms and bonding regions.

At sub-Å resolution, such as the 0.35 Å urea dataset used here, electron density reflects aspherical valence distributions, lone-pair density, and overlap between neighbouring atoms. These chemically structured contributions are preserved under angular averaging. Consequently, the $\ell = 0$ component maintains molecular connectivity and may appear elongated along bond directions rather than spherical when viewed in Cartesian space.

This behaviour is consistent with conventional electron-density maps at comparable resolution, where bonded atoms also exhibit non-spherical density distributions. The observed shape of the $\ell = 0$ component therefore reflects intrinsic bonding structure rather than artefacts of the decomposition method.

#### Clarifications on Mathematical Framework

The following points summarize clarification questions raised during internal review of the Mathematical Framework. Responses are provided to document the mathematical rationale underlying the discrete angular decomposition procedure.

**Question 1: Why does the derivation begin from the conventional Fourier synthesis of electron density?**
**Answer:** The decomposition introduced in this work operates directly on crystallographic structure factors. The natural starting point is therefore the standard Fourier representation of electron density,

$$\rho(\boldsymbol{r}) = \sum_{\boldsymbol{h}} F(\boldsymbol{h})\, \mathrm{e}^{2\pi i\, \boldsymbol{h}\cdot\boldsymbol{r}},$$

which expresses the density as a superposition of reciprocal-lattice plane waves. All subsequent angular projections are applied to this representation, ensuring compatibility with conventional crystallographic Fourier synthesis.

**Question 2: Why are spherical harmonics used for directional decomposition?**
**Answer:**Spherical harmonics form an orthonormal basis on the unit sphere and therefore provide a mathematically natural framework for representing angular structure. Projection onto spherical harmonics allows isotropic and anisotropic components of reciprocal-space density to be separated systematically without modifying the underlying crystallographic model.

**Question 3: Why does Eq. (directional-ifft) contain the complex conjugate $Y^*_{\ell m}$?**
**Answer:** The complex conjugate arises from the standard inner-product definition used in spherical-harmonic analysis. In the continuous formulation, angular coefficients are obtained as

$$f_{\ell m} = \int f(\widehat{\boldsymbol{k}})\, Y^*_{\ell m}(\widehat{\boldsymbol{k}})\, \mathrm{d}\Omega,$$

which ensures orthogonality of spherical-harmonic components. The discrete summation used in this work represents a lattice-sampled approximation to this projection, and therefore retains the conjugate form.

**Question 4: Why is reciprocal space partitioned into radial shells?**
**Answer:** In the continuous spherical formulation, angular decomposition involves integration over spherical shells with volume element

$$k^2\, \mathrm{d}k\, \mathrm{d}\Omega.$$

To approximate this radial integration using discrete reflections, reciprocal space is partitioned into finite shells of thickness $\Delta k$. Grouping reflections into shells provides a discrete representation of spherical integration and ensures that angular contributions are evaluated at approximately constant reciprocal radius.

**Question 5: Why is the shell-weight factor defined as $w(\boldsymbol{h}) = 4\pi \bar{k}_s^2 \Delta k_s / N_s$?**
**Answer:**

In the continuous formulation, integration over angular coordinates yields the radial measure

$$4\pi k^2\, \mathrm{d}k.$$

Discretization into shells therefore produces a finite shell volume

$$\Delta V_s \approx 4\pi \bar{k}_s^2 \Delta k_s.$$

Dividing this volume by the number of observed reflections $N_s$ distributes the shell contribution uniformly among reflections. This produces a discrete Riemann approximation to the continuous spherical integral.

**Question 6: Why divide by the number of observed reflections $N_s$ rather than the number of theoretically possible reflections?**
**Answer:** Division by the number of observed reflections ensures that the shell-weight normalization remains well defined under incomplete reciprocal-space sampling. Using theoretical reflection counts would introduce systematic biases when reflections are missing. The adopted normalization therefore maintains stability across datasets with varying completeness.

**Question 7: Why is the tapering function $W(k)$ included in the shell-weight definition?**

**Answer:** In its simplest form, $W(k) = 1$ and no tapering is applied. Optional tapering functions are introduced only to reduce artefacts arising from finite-resolution truncation. Abrupt termination of reciprocal-space data produces oscillatory artefacts (Gibbs-type ringing) in reconstructed real-space density. Smooth tapering near the resolution limit reduces these effects and improves numerical stability, but is not required for the validity of the method.

**Question 8: Is tapering necessary when high-resolution shells are incomplete?**

**Answer:** Tapering is not strictly required, but may improve stability when outer reciprocal-space shells are sparsely populated. Incomplete high-resolution shells can introduce increased statistical variability in recovered angular components. Gradual attenuation of contributions near $k_{\max}$ reduces sensitivity to such sparsely sampled regions and improves robustness of the angular decomposition.

**Question 9: Why does the method not require explicit spherical Bessel functions?**

**Answer:** In the continuous spherical Fourier–Bessel formulation, radial integrals involve spherical Bessel functions. In the discrete crystallographic setting, these integrals are approximated by shell-wise summation with appropriate volume weighting. This discrete approximation converges to the continuous formulation in the limit of vanishing shell thickness, and therefore avoids explicit valuation of oscillatory Bessel functions.

**Question 10: Why can reflections not simply be summed directly without shell partitioning?**

**Answer:** Direct summation of reflections is sufficient for reconstruction of total electron density. However, angular decomposition requires evaluation of directional components at approximately constant reciprocal radius. Without shell partitioning, angular contributions would be unevenly weighted according to reflection density, leading to biased angular power estimates. Shell-based normalization ensures consistent weighting of reciprocal-space regions.

**Question 11: How does the method behave under incomplete reciprocal-space sampling?**

**Answer:** The shell-weight normalization remains well defined under incomplete sampling because weights are assigned only to observed reflections. Numerical tests involving random and directional removal of reflections demonstrated stable angular behaviour across a range of sampling conditions (Supplementary Section S1). Optional tapering further improves robustness in sparsely sampled outer shells.
